# The epithelial splicing regulator *ESRP2* is epigenetically repressed by DNA hypermethylation in Wilms tumour and acts as a tumour suppressor

**DOI:** 10.1101/2020.11.02.364570

**Authors:** Danny Legge, Ling Li, Whei Moriarty, David Lee, Marianna Szemes, Asef Zahed, Leonidas Panousopoulus, Wan Yun Chung, Yara Aghabi, Jasmin Barratt, Richard Williams, Kathy Pritchard-Jones, Karim T.A. Malik, Sebastian Oltean, Keith W. Brown

**Author notes:** **Corresponding authors:** Keith W. Brown, University of Bristol, School of Cellular and Molecular Medicine Bristol, Biomedical Sciences Building, University Walk, Bristol BS8 1TD, U.K., Sebastian Oltean, Institute of Biomedical & Clinical Sciences, University of Exeter Medical School, St Luke’s Campus, Heavitree Rd, Exeter, EX1 2LU, UK.

## Abstract

Wilms tumour (WT), a childhood kidney cancer with embryonal origins, has been extensively characterised for genetic and epigenetic alterations, but a proportion of WTs still lack identifiable abnormalities. To uncover DNA methylation changes critical for WT pathogenesis, we compared the epigenome of fetal kidney with two WT cell lines, using methyl-CpG immunoprecipitation. We filtered our results to remove common cancer-associated epigenetic changes, and to enrich for genes involved in early kidney development. This identified four candidate genes that were hypermethylated in WT cell lines compared to fetal kidney, of which *ESRP2* (epithelial splicing regulatory protein 2), was the most promising gene for further study. *ESRP2* was commonly repressed by DNA methylation in WT, and this was shown to occur early in WT development (in nephrogenic rests). *ESRP2* expression could be reactivated by DNA methyltransferase inhibition in WT cell lines. When *ESRP2* was overexpressed in WT cell lines, it acted as an inhibitor of cellular proliferation *in vitro,* and *in vivo* it suppressed tumour growth of orthotopic xenografts in nude mice. RNA-seq of the ESRP2-expressing WT cell lines identified several novel splicing targets, in addition to well-characterised targets of ESRP2. We propose a model in which the mesenchymal to epithelial transition that is essential for early kidney development, can be disrupted in to generate WT, either by genetic abnormalities such as *WT1* mutations, or by epigenetic defects, such as *ESRP2* methylation.

## INTRODUCTION

Wilms tumour (WT; nephroblastoma) is an embryonal kidney cancer, affecting approximately 1 in 10,000 children (1, 2). It originates from fetal kidney, due to the failure of the mesenchymal to epithelial transition (MET) that the metanephric blastema undergoes during normal renal development. The metanephric blastema differentiates to form both epithelial and stromal components in the developing kidney. In WT, most tumours retain some of this differentiation potential, giving a so-called “triphasic” histology, with blastemal, epithelial, and stromal elements all being present in varying degrees (3). Premalignant lesions, known as nephrogenic rests (NRs), are associated with many WTs (4). NRs are usually microscopic lesions, seen in the normal kidney adjacent to WTs, and contain undifferentiated fetal kidney cells (4). It is hypothesised that genetic and epigenetic defects occur during renal development that block MET, leading to the formation of NRs, some of which eventually progress to WT (3, 4).

The molecular events underlying WT pathogenesis involve an array of genetic and epigenetic defects (1–3). The earliest genetic mutations associated with WT were found in the *WT1* gene, which was discovered due to its association with several WT-predisposing syndromes. WT1 is essential for normal kidney development, where it plays a critical role in regulating MET (3, 5). Another critical pathway in renal development is the Wnt pathway (3), and mutations in Wnt pathway components such as *CTNNB1* (6, 7) and *WTX (AMER1*) (8) have also been found in WT. Recent genome-wide sequencing studies have identified mutations in microRNA-processing genes, such as *DROSHA, DICER* and *DGCR8,* as well as mutations in other renal developmental regulators, including *SIX1, SIX2* and *SALL1* (9–14). Most of these events do not show a strong association with clinical outcome, but *TP53* mutations are found in the rare anaplastic variant of WT, which has a much poorer prognosis than other subtypes (15).

Epigenetic alterations are also a common feature of WT (16), especially at the 11p15 locus, where the frequent loss of imprinting (LOI) of the fetal growth factor gene *IGF2* (17, 18) is associated with DNA methylation changes at the *H19* differentially methylated region (DMR) (19, 20). We have also reported recurrent LOI at 11p13 involving imprinted *WT1* transcripts (*WT1-AS/AWT1*) (21, 22). Other epigenetic alterations in WT include global hypomethylation (23), DNA hypermethylation at individual tumour suppressor genes such as *RASSF1A* (23–25), and long-range epigenetic silencing of the *PCDHG@* gene clusters (26).

Despite the large number of individual loci identified as bearing genetic and/or epigenetic lesions in WT, a proportion of WTs still lack identifiable driver mutations or epimutations, implying that there are additional novel genes involved in WT pathogenesis (14). We have used genome-wide DNA methylation analysis of WT to identify novel epigenetic lesions, which previously led to the discovery of the first example of long-range epigenetic silencing in WT (26). In this paper we report on further studies in which we have compared two WT cell lines to fetal kidney, to identify additional DNA methylation changes in WT. One of these novel genes, the alternative splicing regulator *ESRP2* (epithelial splicing regulatory protein 2), is known to be important in epithelial to mesenchymal transitions (EMT), and the reverse, MET (27), suggesting that epigenetic deregulation of MET may be an important factor in WT development. We show that *ESRP2* is frequently silenced by DNA hypermethylation in WT, and that it acts as a tumour suppressor gene, regulating alternative splicing in novel genes, some of which affect pathways known to be important in kidney development.

## MATERIALS AND METHODS

### Data availability

MCIP data have been deposited in NCBI’s Gene Expression Omnibus (5) and are accessible through GEO Series accession number GSE153047: https://www.ncbi.nlm.nih.gov/geo/query/acc.cgi?acc=GSE153047. RNA-seq data have been deposited in NCBI’s Gene Expression Omnibus (5) and are accessible through GEO Series accession number GSE154496: https://www.ncbi.nlm.nih.gov/geo/query/acc.cgi?acc=GSE154496.

### Ethical statement

Wilms tumour samples were obtained from Bristol Children’s hospital (BCH), or from collaborators at the Royal Marsden Hospital (RMH), as part of a UK collaboration. Samples were obtained with informed consent (from parent and/or legal guardian for children less than 18 years of age) and with appropriate ethical approval (E5797, South West – Central Bristol Research Ethics Committee (UK)). All methods were performed in accordance with the relevant guidelines and regulations, including those specified in the UK Human Tissue Act 2004. All animal experiments and procedures were approved by the UK Home Office in accordance with the Animals (Scientific Procedures) Act 1986. Mice were maintained at the Biological Services Unit, University of Exeter, UK.

### Cell lines

The Wit49 WT cell line (28) was a kind gift from Prof. Herman Yeger (The Hospital for Sick Children, Ontario, Canada) and 17.94 was derived in the author’s laboratory (29). Cell line identity was confirmed by short tandem repeat (STR) analysis (supplementary figure S1).

WT cell lines and 293 adenovirus-transformed fetal kidney cells (30) were grown in Dulbecco’s modified Eagle’s medium (DMEM) with 10% fetal bovine serum (FBS), 100 U/ml penicillin, 0.1 mg/ml streptomycin and 2mM L-glutamine, at 37°C in 5% CO_2_.

The V200 and E200L cells lines were derived from Wit49 by transfecting 2×10^5^ cells with 200 μg of an inducible expression vector, pBIG2r (31), either empty (V200) or containing an *ESRP2* cDNA insert (E200L). *ESRP2* cDNA was amplified by PCR from IMAGE clone 4810948, using a forward primer containing a BamHI site and a reverse primer containing an EcoRV site plus a FLAG tag (supplementary table S9), then ligated into BamHI/EcoRV-digested pBIG2r (supplementary figure S2A). Only the *ESRP2*-transfected Wit49 cells (E200L) expressed vector-derived *ESRP2* RNA (supplementary figure S2B). Transfected cells were selected and maintained in 50 μg/ml hygromycin B (Santa Cruz Biotechnology). *ESRP2* expression was induced by addition of 2-5 μg/ml doxycycline (Dox, Sigma), with maximum ESRP2 protein expression at 72 to 96 hrs post-induction (supplementary figure S2C).

### Cell growth assays

For mass culture assays, cells were seeded into 6 well plates (1×10^6^ cells per well) and treated with 2 μg/ml Dox or DMSO vehicle control, with medium changes every 3 days.

Cells were trypsinised and counted at days 3, 6 and 8 using a Countess cell counter and trypan blue stain to exclude dead cells.

For colony assays, cells were seeded into 6 well plates (2×10^5^ cells per well) and treated with 2 μg/ml Dox or DMSO vehicle control. Medium was changed every 3 days, then at 14 days cells were fixed, stained with methylene blue and colonies counted manually.

To monitor proliferation in real time, cells were seeded in a 24-well plate at 5×10^3^ cells per well, in quadruplicates, and transferred to the IncuCyte ZOOM live cell imaging system (Essen BioScience). Images were taken in four different fields in each well, every two hours and phase confluence was calculated as a surrogate for growth at each time point.

### Transwell assay

Cells were pre-treated for 4 days with 2 μg/ml Dox or control media, then 2×10^5^ cells were seeded per well (re-suspended in FBS-free DMEM) into a 6 well plate transwell insert (8 μm pore, PET membrane; Falcon, 353093) in a 6-well plate. Wells were filled with 1.7 ml 10% FBS in DMEM to produce a chemotactic gradient. Following 24 hours, inserts were washed and cells on underside of membrane were fixed and stained with crystal violet. Migrated (stained) cells were counted manually using light microscopy.

### Scratch assay

7.5×10^5^ cells were seeded per well in 24 well plates. Cells were treated for 5 days with 2 μg/ml Dox or control media prior to a scratch being performed manually in the centre of the well using a p200 Gilson pipette tip. Wells were washed gently with PBS to remove dead cells from scratch wound site. Control/Dox media was replaced, and wells were analysed at 24 and 48 hours via widefield microscopy and Image J software was used to determine percentage wound closure.

### Cell Trace Violet (CTV) proliferation assay

CTV (ThermoFisher; C34571) was prepared in DMSO and diluted in PBS according to manufacturer’s instructions. 1 ml of diluted CTV stain was used per 1×10^6^ cells. Cells were incubated for 20 minutes at 37°C in the dark before staining was quenched using 10% FBS in DMEM. Stained cells were subsequently seeded into T12.5 flasks. Controls were prepared at this point via fixing 3×10^5^ stained cells in 1% paraformaldehyde. These cells were stored at 4°C, protected from light until the day of the assay. Seeded cells in T12.5 flasks were treated for 6 days with 2 μg/ml Dox or control media. On day 6, intensity of CTV staining was analysed using a Novocyte flow cytometer and FlowJo software.

### 5-Aza-2’-deoxycytidine treatment

Cells were incubated in medium containing 2μM 5-aza-2’-deoxycytidine (azadC; Sigma) for up to 6 days, with a medium change every two days. Control cultures received equivalent volumes of drug solvent (DMSO).

### Xenografting into nude mice

The Wit49 derivatives V200 and E200L were transduced with lentiviral particles expressing firefly luciferase (Amsbio LVP326), then successfully transduced cells were selected with blasticidin, according to the manufacturer’s protocol. Nude male mice two months old (Charles River) were used. General anaesthesia was induced using isoflurane. For orthotopic kidney implantation, an incision was performed in the left flank of the mice, the left kidney was exteriorised and 3×10^6^ cells were injected. Mice were imaged twice weekly using a Xenogen IVIS device, following intraperitoneal injection with luciferin. Once the tumour signal was apparent, mice were injected intraperitoneally with Doxycycline three times per week (50 mg/kg in 5% glucose). Mice were culled either when tumours grew to the maximum allowed size of 10mm in diameter (according to the animal licence), or after two months of imaging. Dissected tumours were analysed by Western Blot for the induction of ESRP2.

### DNA extraction and methyl CpG immunoprecipitation (MCIP)

DNA was extracted from WT cell lines with a DNeasy kit (Qiagen). Human fetal kidney DNA was obtained from BioChain. MCIP was performed as described previously (32) by co-hybridising methylation-enriched DNA fractions with input DNA on to a custom microarray (Nimblegen), based on design 2006-04-28_HG18_Refseq_Promoter (see GEO entry for further details). Statistical analyses by ChIPMonk software (http://www.bioinformatics.bbsrc.ac.uk/projects/chipmonk), used windowed T-tests to identify differentially methylated genes (supplementary table S1). log2 gene methylation levels were derived from the mean probe ratios within 700bp of the transcriptional start site. The MCIP data have been deposited in NCBI’s Gene Expression Omnibus (5) and are accessible through GEO Series accession number GSE153047: https://www.ncbi.nlm.nih.gov/geo/query/acc.cgi?acc=GSE153047.

### Pyrosequencing

DNA was purified by phenol-chloroform extraction, bisulfite converted (EZ DNA Methylation Gold kit; Zymo Research), amplified using a Pyromark PCR kit (Qiagen) and pyrosequenced on a PyroMark Q96 instrument (Qiagen), using primers listed in supplementary table S10.

### RNA extraction, cDNA synthesis and RT-PCR

Total RNA was extracted using TriReagent (Sigma) and DNase treated with TURBO DNA-free (Ambion). Human fetal kidney RNA was obtained from BioChain. cDNA was synthesized using the Superscript IV RT-PCR system (Invitrogen). Gene-specific primers (supplementary table S9) were used for end-point PCR (HotStarTaq Plus DNA Polymerase; Qiagen), to detect different sized amplicons on agarose gels (1.5%), representing inclusion or exclusion of alternative exons. Quantitative real-time PCR (QPCR) using gene-specific primers (supplementary table S9) was performed using QuantiNova SYBR Green mix (Qiagen) on an MX3000P real-time PCR machine (Stratagene), normalising the amount of target gene to the endogenous level of *TBP.* Human universal RNA (Agilent) was used as a reference to standardise results between QPCR batches.

### Protein extraction and Western Blotting

Cells were washed with ice-cold PBS and lysed in cell lysis buffer (Cell Signaling), with complete mini inhibitors (Roche) for 10 min on ice, and then sonicated for 5 min at high intermittent pulses (30/30) (Diagenode, Bioruptor). Samples were centrifuged for 10 min at 10,000 g at 4°C to remove any cell debris and typically 25 μg proteins were separated on SDS-polyacrylamide gels and analysed by Western blotting. Primary antibodies were against ESRP2 (rabbit, Abcam ab155227), FLAG (mouse, Sigma F3165) and β-ACTIN (rabbit, Abcam AB8227), followed by secondary HRP-labelled anti-rabbit IgG (Sigma A6154) or anti-mouse IgG (Sigma A9044). Chemiluminescence detection was with Lumiglo (KPL) and X-ray films were imaged on a flatbed scanner and analysed using Image J (http://imagej.nih.gov/ij/).

### Immunofluorescence

Cells were grown on sterile glass slides, fixed for 30 min at room temperature in 1% paraformaldehyde in phosphate buffered saline (PBS), permeabilised for 10 min in 0.5% Triton X-100 in PBS and finally rinsed in 50mM glycine in PBS. Fixed cells were then stained by indirect immunofluorescence using a primary antibody against FLAG (mouse, Sigma F3165) and secondary antibody against mouse IgG (Alexa Fluor 488-labelled; Invitrogen) to detect the transfected ESRP2, together with Alexa Fluor 594-labelled phalloidin (Invitrogen) to detect actin. Antibodies were diluted in PBS +1% bovine serum albumin, containing 0.1 μg/ml DAPI to image nuclei. Slides were mounted in Fluoroshield (Sigma) and examined with a confocal microscope, acquiring eight images at 1μm spacing per field. Maximum intensity projections were merged using Image J software (http://imagej.nih.gov/ij/).

### RNA sequencing (RNA-seq)

RNA was extracted from E200L cells 96 hrs after treatment with 2 μg/ml Dox, or control solvent (DMSO), using an RNAeasy kit (Qiagen), then DNase treated, and quality confirmed using an Agilent Screen tape RNA assay. Two biological replicates were used for RNA-seq (i.e. four samples total). Sequencing libraries were prepared from total RNA (500 ng) using the TruSeq Stranded mRNA Library Preparation kit (Illumina, Inc.) and uniquely barcoded adapters (RNA LT adapters, Illumina, Inc). Libraries were pooled equimolarly for sequencing, which was carried out on the NextSeq500 instrument (Illumina, Inc.) using the NextSeq High Output v2 150-cycle kit (Illumina, Inc.). Approximately 300 million paired reads (passing filter, PF) were obtained, divided between the four experimental samples. NextSeq Control Software Version 2.0.0 and RTA v2.4.6 were used for instrument control and primary analysis respectively. Reads from the four samples were mapped to the human genome (hg19) using the new Tuxedo Suite of programs (HISAT2, StringTie, Ballgown; https://www.ncbi.nlm.nih.gov/pubmed/?term=27560171). To identify RNA splicing alterations, the four BAM files generated by HISAT2 were used as input for rMATS ((33), replicate Multivariate Analysis of Transcript Splicing; http://rnaseq-mats.sourceforge.net/user_guide.htm). Bam files were viewed in the Integrative Genomics Viewer (IGV; http://software.broadinstitute.org/software/igv/) to produce Sashimi plots of alternative splicing. The RNA-seq data have been deposited in NCBI’s Gene Expression Omnibus (5) and are accessible through GEO Series accession number GSE154496: https://www.ncbi.nlm.nih.gov/geo/query/acc.cgi?acc=GSE154496.

## RESULTS

### Genome-wide DNA methylation analysis

To identify novel epigenetically-regulated genes involved in WT pathogenesis, methyl CpG immunoprecipitation (MCIP) and promoter microarrays were used to detect regions that were DNA methylated in human fetal kidney and in two WT cell lines; Wit49 (28) and 17.94 (29). Windowed t-tests in ChIPMonk software were used to identify 225 genes that were hypermethylated in the WT cell lines compared with fetal kidney (figure 1A and supplementary table S1). Gene ontology analysis showed that this set of genes was especially enriched in genes involved in chromatin organisation, developmental processes, and transcriptional regulation (figure 1B and supplementary table S2).

**Figure 1:**
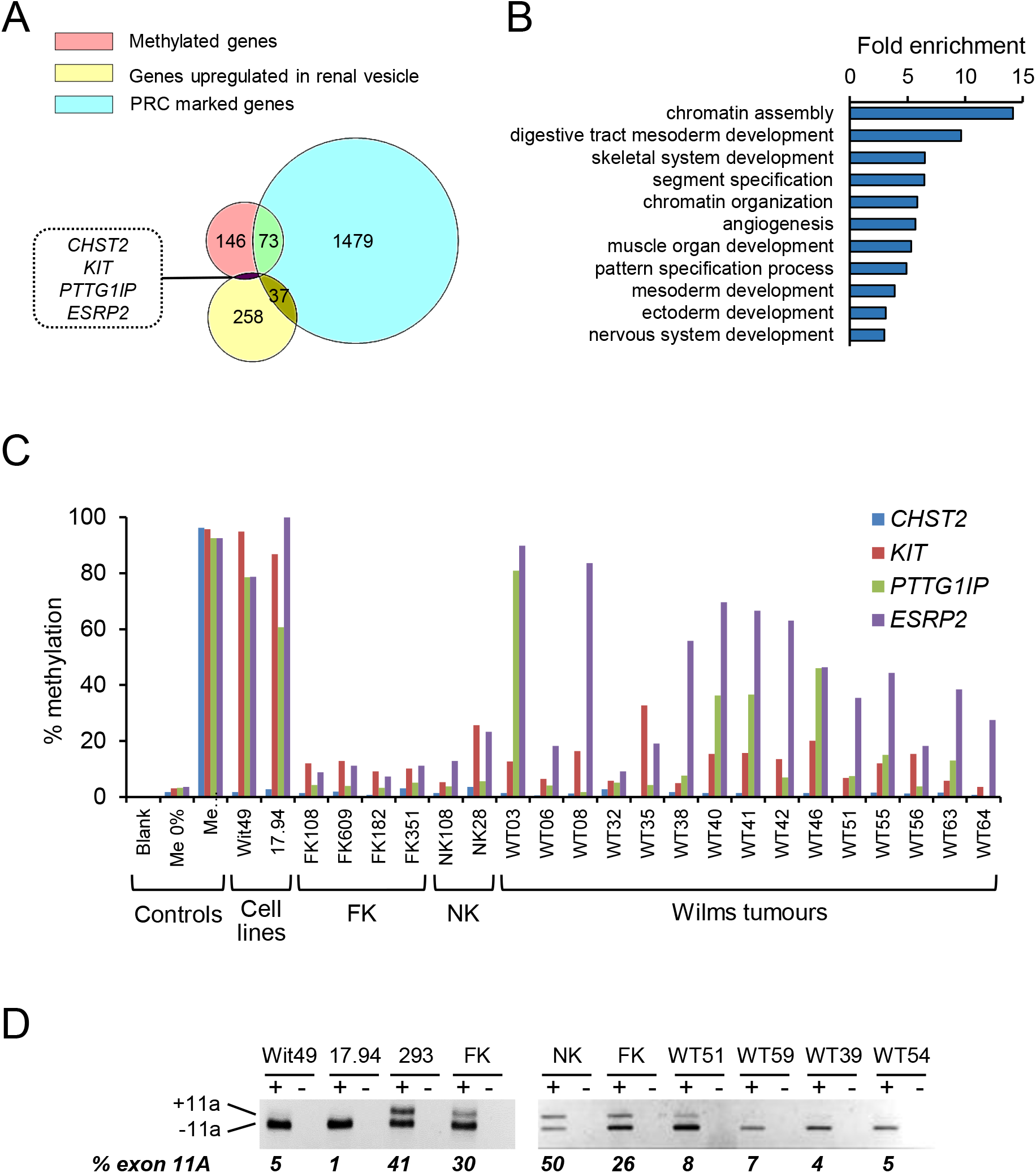
Identification of *ESRP2* as a candidate hypermethylated gene. A: Venn diagram showing filtering of the 225 methylated genes that were identified by MCIP, firstly by negative selection for genes that are PRC marked in ES cells (34) and secondly by positive selection for genes that are upregulated in the renal vesicle during kidney development (38). The full list of methylated genes and filtered lists are shown in supplementary table S1. B: Gene ontology (GO) analysis of the 225 methylated genes. Only categories with a fold enrichment >3 are shown; see supplementary table S2 for full results. C: The four candidate genes were analysed for DNA methylation by pyrosequencing in two WT cell lines, four FK, two NK and 15 WT. Me 0% and Me 100% are unmethylated and fully methylated DNA controls. See supplementary table S10 for pyrosequencing primers. D: Alternative splicing of *ENAH* exon 11A was analysed by RT-PCR followed by agarose gel electrophoresis to detect different sized amplicons, in two WT cell lines (Wit49 and 17.94), normal fetal kidney cell line 293, FK, NK and four WTs.

To define a group of genes that were specifically methylated in WT, two filters were applied to the list of methylated genes. Firstly, genes were removed that are polycomb repressive complex (PRC) marked in embryonic stem (ES) cells (34), since such genes seem predisposed to become DNA methylated in a wide range of human cancers (35–37) and therefore would probably not be WT-specific. Secondly, a positive selection was applied for genes whose expression is upregulated in the renal vesicle early in kidney development (38), reasoning that epigenetic inactivation of such genes could be involved in the block in mesenchymal to epithelial transition (MET) that is critical for WT development (3). Using these criteria, four candidate genes were identified: *CHST2, KIT, PTTG1IP* and *ESRP2 (RBM35B*) (figure 1A and supplementary table S1). Pyrosequencing was then used to examine the DNA methylation levels of these four genes in fetal kidney, normal kidney and a small number of Wilms tumours. Of the four candidate genes, *ESRP2* was the most consistently methylated in WT (figure 1C).

*ESRP2* was a particularly attractive candidate gene for further study, because of its previously described involvement in the control of alternative splicing events necessary for epithelial cell differentiation, which are abrogated in EMT in cancer (39). Supporting evidence for a possible role for *ESRP2* inactivation in WT came from examination of the *ESRP2* splicing target *ENAH.* ESRP2 induces the inclusion of the epithelial-specific alternatively spliced exon 11a in *ENAH* RNA transcripts (40). Using RT-PCR across the alternatively spliced exon, less of the epithelial isoform of *ENAH* (+exon 11A) was expressed in WTs compared to normal kidney (NK) and fetal kidney (FK), consistent with down-regulation of *ESRP2* in WT (figure 1D). We therefore went on to examine DNA methylation and expression of *ESRP2* in two large cohorts of WTs.

### DNA methylation of ESRP2 in Wilms tumour

A pyrosequencing assay was used to examine DNA methylation in the *ESRP2* promoter CpG island (CGI); this assay was located centrally in the region that was identified by MCIP as being hypermethylated in the WT cell lines compared to fetal kidney (supplementary figure S3). In one cohort of WTs, the methylation of the closely related paralog *ESRP1* was also studied by pyrosequencing.

The first cohort of WTs was from Bristol Children’s Hospital (BCH) and consisted of tumour samples of all stages, mostly obtained at surgical resection, pre-chemotherapy. Of this group, 72% of WTs were hypermethylated (defined as DNA methylation >25% at the *ESRP2* promoter) compared to normal kidney tissue (NT) (figure 2A and supplementary figure S4A; NT mean=13%, 95% CI=9-17%; WT mean=44%, 95% CI=38-50%; means differ by 31%; p<0.0005, t-test).

**Figure 2:**
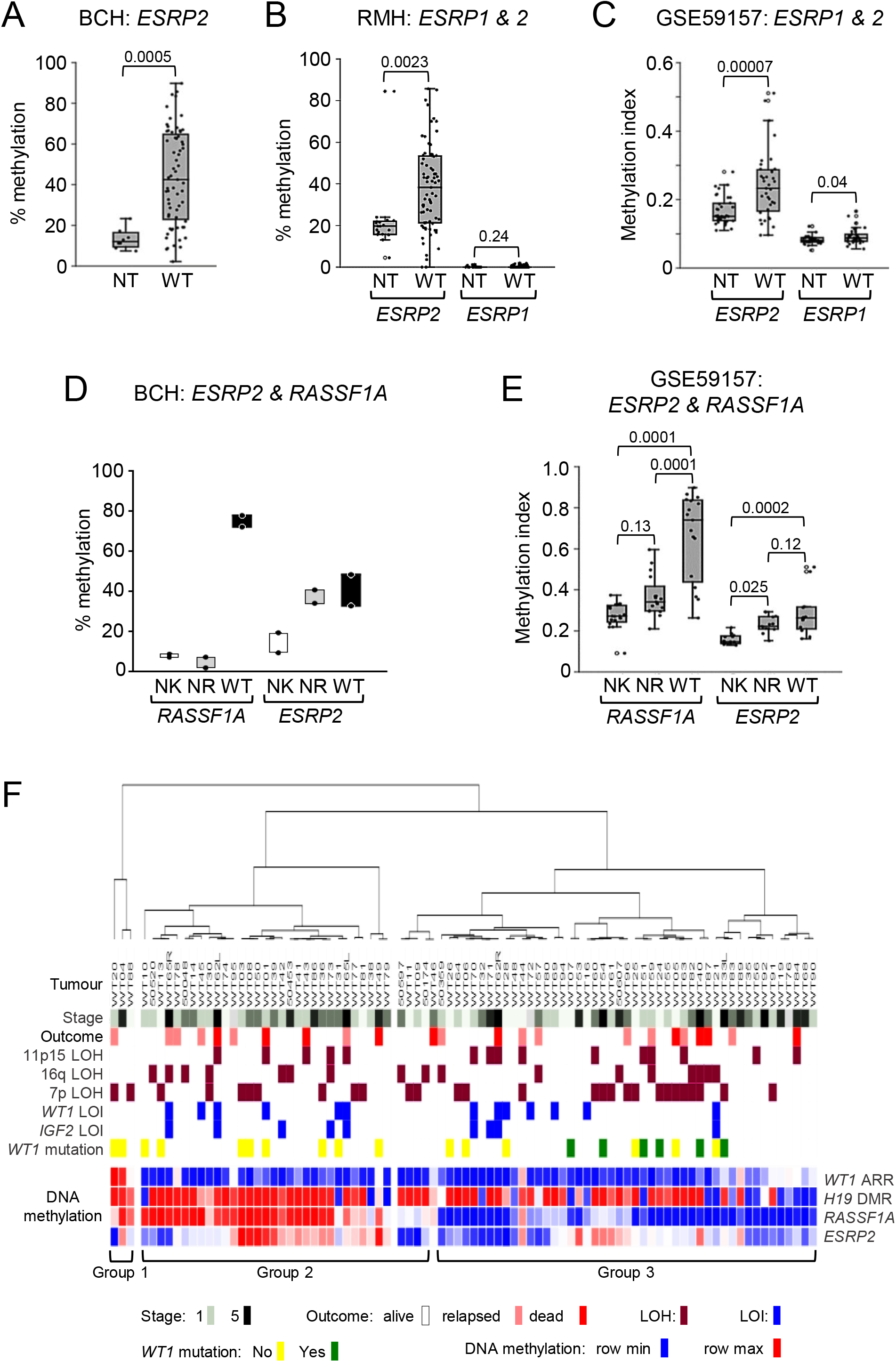
*ESRP2* is hypermethylated in Wilms tumours and nephrogenic rests. A: Dot-boxplot of *ESRP2* DNA methylation in the BCH cohort. NT n=8 (4 NK and 4 FK), WT n=65, p value from t-test. B: Dot-boxplot of *ESRP2* and *ESRP1* DNA methylation in the RMH cohort. *ESRP2:* NT n=18 (all NK), WT n=73; *ESRP1:* NT n=15 (all NK), WT n=69, p values from t-test. C: Dot-boxplot of *ESRP2* and *ESRP1* DNA methylation in dataset GSE59157. NT n=36 (all NK), WT n=37, p values from t-test. D: *RASSF1A* and *ESRP2* methylation in nephrogenic rests in the BCH cohort. Two sets of matched NK, NR and WT are shown for *RASSF1A* and *ESRP2* methylation. E: Dot-boxplot of *RASSF1A* and *ESRP2* methylation in nephrogenic rests in the GSE59157 dataset. 17 sets of matched NK, NR and WT are shown for *RASSF1A* methylation and 13 sets for *ESRP2* methylation, from individuals where the WT was hypermethylated compared to the matched NK. p values from Tukey’s pairwise test. F: Hierarchical clustering of Wilms tumours by DNA methylation. WTs from the BCH cohort were assayed for DNA methylation at the *WT1* antisense regulatory region (ARR), the *H19* differentially methylated region (DMR), the *RASSF1A* promoter and the *ESRP2* promoter. Details of the pyrosequencing assays are given in supplementary table S10 and full methylation results in supplementary table S3. Loss of heterozygosity (LOH), loss of imprinting (LOI), stage and survival data taken from refs (21, 45), and *WT1* mutation data from refs (47, 48, 64, 84). Samples were clustered by one minus Pearson correlation using Morpheus software: https://software.broadinstitute.org/morpheus/Methylation. DNA methylation was assayed by pyrosequencing in A, B, D and F, and by Illumina Human Methylation 450K BeadChips in C and E.

The second cohort was from samples collected at the Royal Marsden Hospital (RMH) as part of a UK-wide trial. These samples were from stages 1 to 3 and were taken at surgical resection, post-chemotherapy. In this different cohort, 78% of WTs were hypermethylated (figure 2B and supplementary figure S4C; NT mean=22%, 95% CI=14-30%; WT mean=38%, 95% CI=33-44%; means differ by 16%; p=0.0023, t-test). *ESRP1* DNA methylation was also tested in the RMH cohort and found to be very low (<2%) in both NT and WT, with no significant difference between NT and WT methylation (figure 2B; p=0.24, t-test).

Additional independent DNA methylation data was extracted for the *ESRP1* and *ESRP2* promoter CGIs from the publicly available dataset GSE59157 (figure 2C), which examined WTs using Illumina bead chips (41). This dataset also showed hypermethylation of *ESRP2* in WTs (NT mean=0.17, 95% CI=0.15-0.18; WT mean=0.24, 95% CI=0.21-0.27; means differ by 0.08; p=0.0007, t-test), with much lower methylation of *ESRP1,* that was only marginally different in NT and WT (NT mean=0.08, 95% CI=0.08-0.09; WT mean=0.09, 95% CI=0.09-0.10; means differ by 0.01; p=0.04, t-test).

Thus, *ESRP2* DNA was found to be hypermethylated in two independent cohorts of WTs and these findings were confirmed in another independent set of publicly available data. However, the *ESRP2* paralog *ESRP1* was not found to be hypermethylated in WT, despite the similar biological functions described for the two paralogs (42).

The relationship between tumour stage and *ESRP2* DNA methylation was investigated in the two cohorts of tumours but there was no significant association with stage in either cohort (supplementary figures S5A and B). WTs from both cohorts were divided into hypermethylated (>25% methylation) and non-methylated groups (<25% methylation), and overall patient survival examined. There was no association between *ESRP2* methylation and survival, with opposite non-significant trends in the two cohorts (supplementary figure S6). These results suggest that *ESRP2* methylation is not associated with different clinical stages or outcomes in WT.

*ESRP2* is located on chromosome 16q22 (https://www.ncbi.nlm.nih.gov/gene/80004), which is a chromosomal region found to show frequent loss of heterozygosity (LOH) in WT (43).

The relationship between 16q LOH and *ESRP2* DNA methylation was therefore tested in the BCH cohort, which had previously been characterised for 16q LOH (21). No difference was observed in the *ESRP2* methylation in WTs with or without 16q LOH (supplementary figure S5C).

Most WTs are thought to develop via premalignant lesions, known as a nephrogenic rests (NRs), which are frequently found in the kidney tissue adjacent to WTs (4). To characterise the phase of WT development at which *ESRP2* DNA methylation occurs, methylation was assayed in two sets of matched NK, NR, and WT from the BCH cohort. *ESRP2* was DNA hypermethylated at a similar level in NR and the matched WT compared to NK (figure 2D). In contrast, *RASSF1A,* a tumour suppressor gene that is frequently hypermethylated in WT (24), was not hypermethylated in NRs (figure 2D), as previously reported (23). Dataset GSE59157 (discussed above) also contains methylation data on nephrogenic rests, so methylation values for the *ESRP2* and *RASSFIA* promoter CGIs were extracted for all WTs that showed hypermethylation in these genes compared to their matched NK samples (figure 2E). In those tumours, *ESRP2* was significantly more methylated in WT compared to NK (NK mean=0.16, 95% CI=0.14-0.17; WT mean=0.28, 95% CI=0.22-0.35; means differ by 0.12; p=0.0002, Tukey’s pairwise test), but there was no significant difference between WT and NR methylation (NR mean=0.23, 95% CI=0.21-0.26; WT mean=0.28, 95% CI=0.22-0.35; means differ by 0.05; p=0.12, Tukey’s pairwise test) but NRs were hypermethylated compared to NK (NK mean=0.16, 95% CI=0.14-0.17; NR mean=0.23, 95% CI=0.21-0.26; means differ by 0.07; p=0.025, Tukey’s pairwise test). In contrast, whilst *RASSF1A* was hypermethylated in WTs compared to NK (NK mean=0.27, 95% CI=0.24-0.31; WT mean=0.66, 95% CI=0.56-0.77; means differ by 0.39; p=0.0001, Tukey’s pairwise test), NRs were not significantly hypermethylated compared to NK (NK mean=0.27, 95% CI=0.24-0.31; NR mean=0.37, 95% CI=0.32-0.42; means differ by 0.10; p=0.12, Tukey’s pairwise test). Therefore, in both the BCH and GSE59157 datasets, *ESRP2* appeared to be hypermethylated in NRs compared to NK, whereas *RASSF1A* was not, implying that *ESRP2* hypermethylation is an earlier event in WT development compared to *RASSF1A* hypermethylation.

To investigate whether epigenetic changes, including *ESRP2* hypermethylation in WT, are associated with other clinical and molecular features, the BCH cohort of WTs were studied, since these WTs have been extensively investigated for LOH, loss of imprinting (LOI), *WT1* mutations and their relationship to clinical parameters (21, 22, 44–50). In addition to *ESRP2* methylation, DNA methylation was measured by pyrosequencing at other loci previously shown to have consistent epigenetic changes in WT; the *WT1* antisense regulatory region (ARR) (22), the *H19* DMR (21) and *RASSF1A* (24). WTs were then grouped by hierarchical clustering of the DNA methylation values at the four loci (figure 2F and supplementary table S3), which divided them into two major clusters and one minor cluster. There were no associations between these clusters and either stage, outcome, LOH or LOI (figure 2F and supplementary table S3). Interestingly, all tumours with *WT1* mutations were in the same cluster (group 3), in which there was less methylation of *RASSF1A* and *ESRP2* compared to the other two groups (figure 2F and supplementary table S3). This may suggest that *WT1* mutation and DNA hypermethylation of certain genes, including *ESRP2,* are mutually exclusive.

*ESRP2* DNA methylation was also examined in 16 non-WT childhood renal tumours from BCH (supplementary figure S7A) and hypermethylation was observed in 10 (63%), which was most marked in clear cell sarcomas of the kidney (CCSK) and in rhabdoid tumours (RH). In two publicly available datasets (GSE73187 (51) and GSE4487 (52)), *ESRP2* was also found to be hypermethylated in CCSK (supplementary figures S7B and C) and in RH (supplementary figure S7C), suggesting that epigenetic inactivation of *ESRP2* may be involved in the pathogenesis of several types of renal tumours of childhood, not just WT.

### Expression of ESRP2 in Wilms tumour

To determine whether the DNA hypermethylation of *ESRP2* leads to epigenetic deregulation of *ESRP2* expression, *ESRP2* RNA expression was analysed by QPCR in the two WT cohorts (figure 3 and supplementary figure S4).

**Figure 3:**
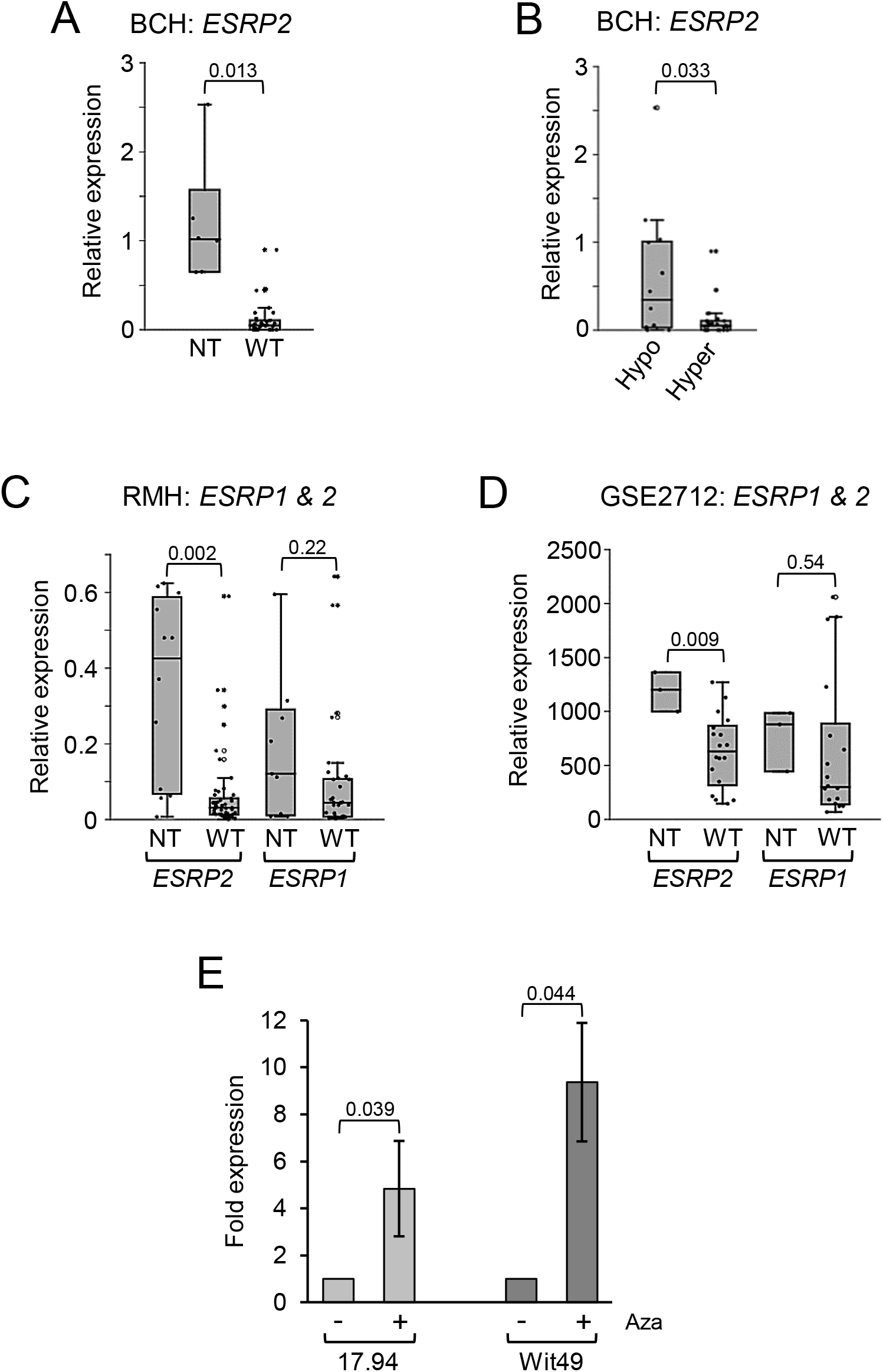
*ESRP2* expression is repressed in Wilms tumour and regulated by DNA methylation. A: Dot-boxplot of *ESRP2* RNA expression relative to FK in the BCH cohort. NT n=6 (3 NK and 3 FK), WT n=32, p value from t-test. B: Dot-boxplot of *ESRP2* expression in hypomethylated versus hypermethylated samples in the BCH. Hypomethylated n=14 (*ESRP2* methylation<25%, Hypo), Hypermethylated n=24 (*ESRP2* methylation>25%, Hyper), p value from t-test. C: Dot-boxplot of *ESRP2* and *ESRP1* RNA expression relative to NT in the RMH cohort. *ESRP2:* NT n=12 (all NK), WT n=51; *ESRP1:* NT n=9 (all NK), WT n=33, p values from t-tests. D: Dot-boxplot of *ESRP2* and *ESRP1* RNA expression in the GSE2712 dataset. NT n=3 (all FK), WT n=18, p values from t-tests. E: 17.94 and Wit49 WT cell lines were treated with 2μM 5-aza-2’-deoxycytidine (Aza) for six days. *ESRP2* RNA expression expressed relative to untreated cells. Results are mean ± SEM of n=3, p values from paired t-tests. RNA expression was measured by real-time QPCR, normalized to endogenous levels of *TBP* in A, B, C and E, and by Affymetrix Human Genome U133A Array in D.

In the BCH cohort, expression of *ESRP2* in WT was very low compared to normal kidney tissue (figure 3A and supplementary figure S4B) (NT mean=1.2, 95% CI=0.5-1.9; WT mean=0.1; 95% CI=0.0-0.2; means differ by1.1; p=0.013, t-test). When WTs were divided into two groups by their *ESRP2* methylation status (figure 3B), hypermethylation was seen to be associated with reduced expression of *ESRP2* (hypomethylated mean=0.57; 95% CI=0.17-0.97; hypermethylated mean=0.11; 95% CI=0.03-0.19; means differ by 0.46; p=0.033, t-test).

In the RMH cohort, expression of *ESRP2* was also reduced in WT compared to NT (figure 3C and supplementary figure S4D) (NT mean=0.35; 95% CI=0.20-0.50; WT mean=0.06; 95% CI=0.03-0.09; means differ by 0.29; p=0.002, t-test). *ESRP1* expression was also examined in the RMH cohort (figure 3C) but it was not significantly different in WT compared to NT (NT mean=0.18, 95% CI=0.04-0.32; WT mean=0.09, 95% CI=0.04-0.15; means differ by 0.09; p=0.22, t-test).

These findings were confirmed using publicly available microarray data (GSE2712;(53)) on gene expression in WTs versus NT (figure 3D). In this dataset *ESRP2* expression was lower in WT compared to NT (NT mean=1188, 95% CI=734-1641; WT mean=631, 95% CI=464-798; means differ by 556; p=0.009, t-test), but *ESRP1* expression did not differ significantly (NT mean=770, 95% CI=56-1483; WT mean=622, 95% CI=290-954; difference between means=148, p=0.54, t-test).

These results showed that *ESRP2* but not *ESRP1* expression was reduced in WTs compared to NT and that reduced expression of *ESRP2* was associated with hypermethylation. This suggested a mechanistic link between *ESRP2* methylation and gene expression, so this was tested directly using a pharmacological inhibitor of DNA methylation on two hypermethylated WT cells lines (methylation data shown in figure 1C and supplementary figure S4A). When the two WT cell lines were treated with the DNA methylation inhibitor 5-aza-2’-deoxycytidine (Aza), there was a 5 to 10-fold increase in *ESRP2* RNA expression (p<0.05, paired t-test) (figure 3E).

### Biological function of ESRP2 in vitro

The methylation and expression results described above suggested that *ESRP2* may have an important functional role in the development of WT. To carry out functional analyses on *ESRP2* in Wilms tumour, the WT cell line Wit49 was transfected with an expression vector that encodes full-length human *ESRP2* (with a FLAG tag) under the control of a doxycycline (Dox)-inducible promoter (supplementary figure S2). The cell line (E200L) showed strong Dox-induced expression of *ESRP2* RNA (figure 4A) and protein (figure 4B), with the expected nuclear localisation of ESRP2 protein (figures 4F and G). Induced expression of ESRP2 drove the splicing of the known target gene *ENAH* (40) towards its epithelial splice form, with increased expression of transcripts containing exon 11a (figure 4C), demonstrating that the inserted ESRP2 construct produced biologically active protein in the WT cell line.

**Figure 4:**
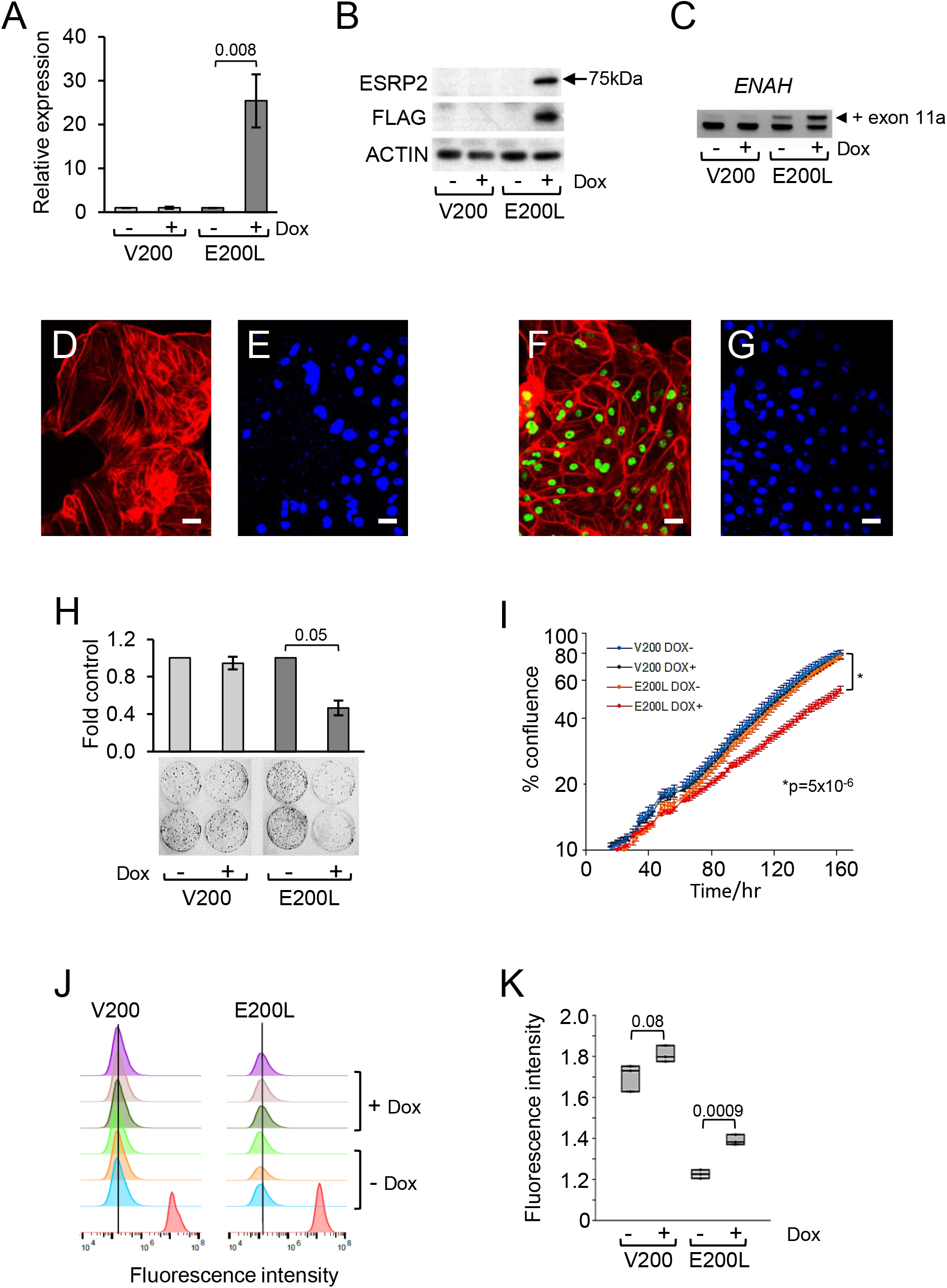
Inducible expression of *ESRP2* in a Wilms tumour cell line. A: *ESRP2* RNA expression assayed by QPCR, normalized to endogenous levels of *TBP,* in V200 and E200L cells after 72 hr Dox induction, shown as fold induction relative to uninduced cells. Results are mean ± SEM of n=3, p value from paired t test. B: ESRP2 protein assayed by Western blotting in V200 and E200L cells after 72 hr Dox induction. Anti-ESRP2 detected total ESRP2 protein, anti-FLAG detected vector-derived ESRP2 and anti-ACTIN was used as a loading control. Representative of three experiments. C: Alternative splicing of *ENAH* exon 11A was analysed by RT-PCR followed by agarose gel electrophoresis to detect different sized amplicons, in V200 and E200L cells after 72 hr Dox induction. Representative of three experiments. D to G: Immunofluorescence of E200L cells, stained for FLAG-tagged ESRP2 (green) and ACTIN (red) after 72 hr Dox induction (F), or uninduced (D). E and G are DAPI staining of the nuclei in the same fields shown in D and F. Scale bars = 50 μm. H: Colony-forming assay of induced (Dox-treated) and uninduced V200 and E200L cells, shown as fold colony numbers compared to uninduced controls after 14 days. Results are mean ± SEM of n=3, p values from paired t test. I: Cell confluence assay (by IncuCyte), showing growth of induced (Dox-treated) and uninduced V200 and E200L cells. Results are mean ± SEM of n=3, p value at 162 hrs from paired t test. Representative of three independent experiments. J, K: Cell Trace Violet (CTV) proliferation assay of induced (Dox-treated) and uninduced V200 and E200L cells. J: CTV staining of triplicates of induced and uninduced cells showing median fluorescence intensity histograms at 6 days of treatment. Red peaks are controls representing staining of cells at day zero. K: Dot-boxplot of quantitation of staining at 6 days (n=3). p values from paired t tests of log-transformed values.

ESRP2 expression affected biological properties of the transfected cells. Over-expression of ESRP2 was associated with an apparent redistribution of actin filaments towards the cell periphery (figure 4F, G), compared to the more cytoplasmic distribution of actin stress fibres seen in the uninduced cells (figure 4D, E). This is like observations of the effect of overexpression of ESRPs in other systems (54, 55).

Dox-induced ESRP2 overexpression caused decreased colony-forming efficiency in the E200L WT cell line (figure 5H), as well as a reduced growth rate in mass cultures (supplementary figure S8). Real-time analysis of cell density showed a slower cell proliferation rate in the Dox-induced E200L cells (figure 4I), which was associated with a decreased rate of cell division as assessed by Cell Trace Violet (CTV) staining (figure 4J, K). Measurement of cell invasion using a transwell assay (supplementary figure S9A) and of cell motility using a scratch assay (supplementary figure S9B), showed no differences between EPRP2-induced and uninduced WT cells. These results therefore suggested that downregulation of ESRP2 expression may play a role in loss of growth control in Wilms tumour cells, but probably not in cell motility and invasion.

**Figure 5:**
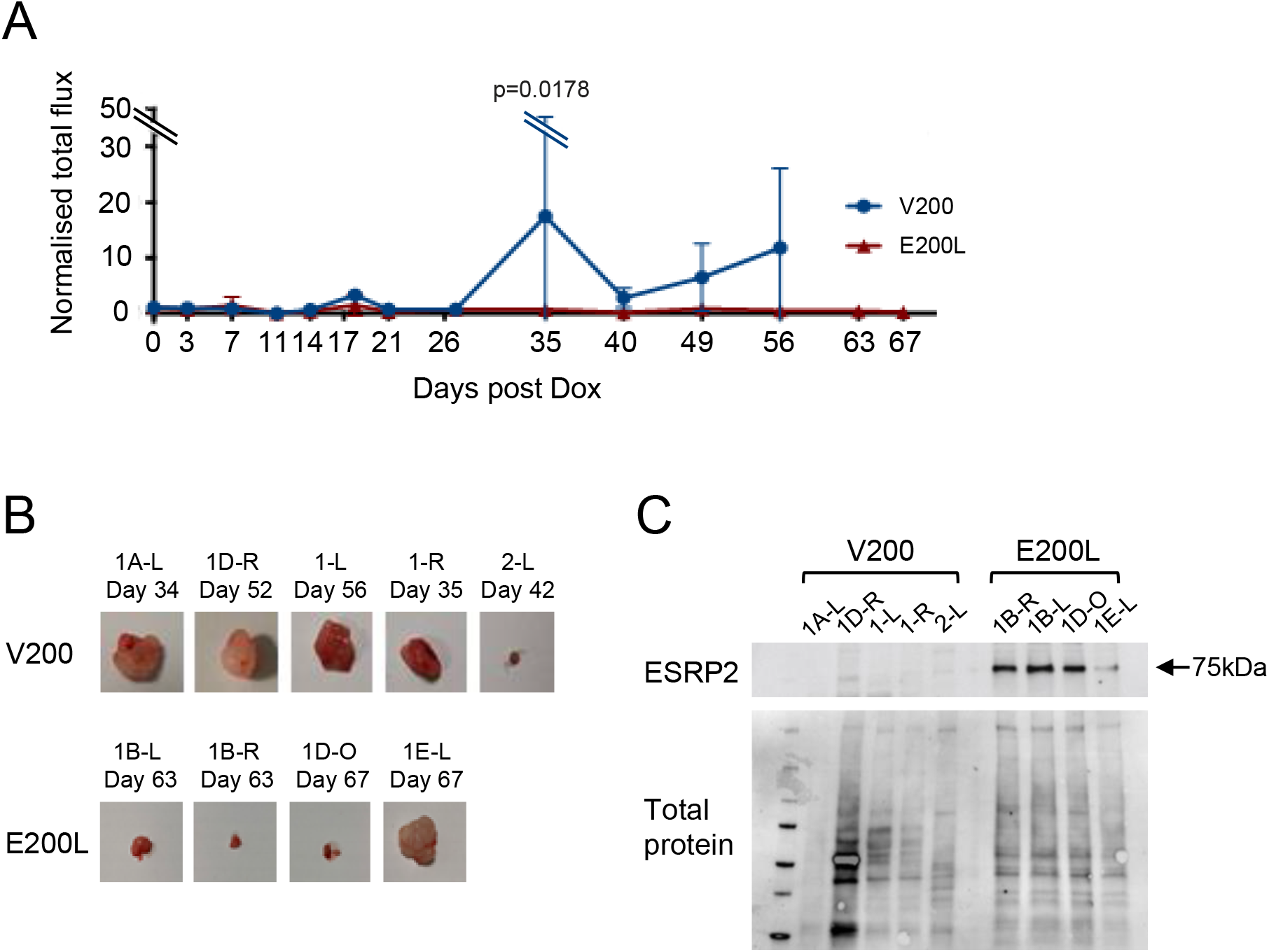
Tumorigenicity of ESRP2-expressing WT cells. Orthotopic xenografts of V200 and E200L cells we produced by injecting cells under the kidney capsule of nude mice (n=5 for each cell line). When a bioluminescent signal was detected, mice were injected intraperitoneally with Dox three times per week, as described in Materials and Methods. A: Time course of tumour growth as assayed by *in vivo* bioluminescence in V200 and E200L xenografts. Plot shows the average ± SEM of tumour signals normalized to initial signal (i.e., the start of Dox induction). p=0.0178 by two-way ANOVA. Full traces of individual tumour growth and examples of bioluminescence are shown in supplementary figure S10A to D. B: Tumours excised from mice injected with V200 and E200L cells (no tumour was excisable in mouse 1B-0 at day 63, therefore only four E200L tumours are shown). Full details of tumour size and weight are shown in supplementary figure S10E. C: Western blot of ESRP2 protein expression in tumours excised from V200 and E200L implanted mice.

### Xenograft assays of ESRP2 function in vivo

The effect of ESRP2 expression on tumorigenicity *in vivo*, was tested by injecting the Wit49-derived cell lines into nude mice (figure 5). We used orthotopic xenografts under the kidney capsule, since this procedure has been shown to give a more authentic WT-like histology than subcutaneous xenografts (56). Monitoring of tumour growth by whole-body luciferase fluorescence imaging showed that after treatment with Dox, tumours produced by injecting V200 cells continued to proliferate, whilst tumours produced by E200L cells stopped growing, or regressed (figure 5A and supplementary figure S10 A to D). Injection of V200 cells produced large tumours in four of five mice, but injection of E200L cells only produced a large tumour in one mouse out of five (1E-L) (figure 5B and supplementary figure S10E). Interestingly, Western blotting of excised tumours demonstrated that Dox treatment of the mice had induced high-level ESRP2 expression in all E200L tumours, with the notable exception of 1E-L (where the tumour grew larger) and V200-induced tumours (figure 5C), demonstrating a strong correlation between ESRP2 expression and suppression of tumour growth. Thus, ESRP2 expression in the Wit49 cell line acts as a tumour suppressor in an orthotopic model of *in vivo* tumorigenicity.

### RNA-seq analysis of alternative splicing in Wilms tumour cell lines

The evidence just described suggested that ESRP2 is epigenetically repressed in Wilms tumours, where it can act as functional tumour suppressor. Given the previous identification of ESRP2 as a regulator of alternative splicing (27, 39, 42), we used RNA sequencing (RNA-seq) of the Wit49-derived cell lines to identify alternative splicing events that could contribute to the pathogenesis of Wilms tumour. In biological duplicates, we extracted RNA from E200L cells that were Dox-induced (ESRP2-expressing) or uninduced (non-expressing), and sequenced them, obtaining between 70 and 80 million paired-end reads per sample. These reads were mapped onto the human genome, examined for differential gene expression, and used in rMATS software (33) to identify alternative splicing events.

Very few transcripts, apart from *ESRP2,* showed significant changes in RNA expression (p<0.05, fold change >2) when ESRP2 expression was induced in E200L cells (figure 6A and B). Interestingly, one induced gene was *GRHL1,* and the grainyhead-like transcription factors have been shown to be important in both kidney development and MET (57), making them good candidates for an involvement in WT. Using a specific QPCR assay, we confirmed the induction of *GRHL1* expression following ESRP2 induction (supplementary figure S11A), but we found no difference in expression of *GRHL1* between NT and WT (supplementary figure S11B), which does not support a role for altered *GRHL1* expression in WT pathogenesis.

**Figure 6:**
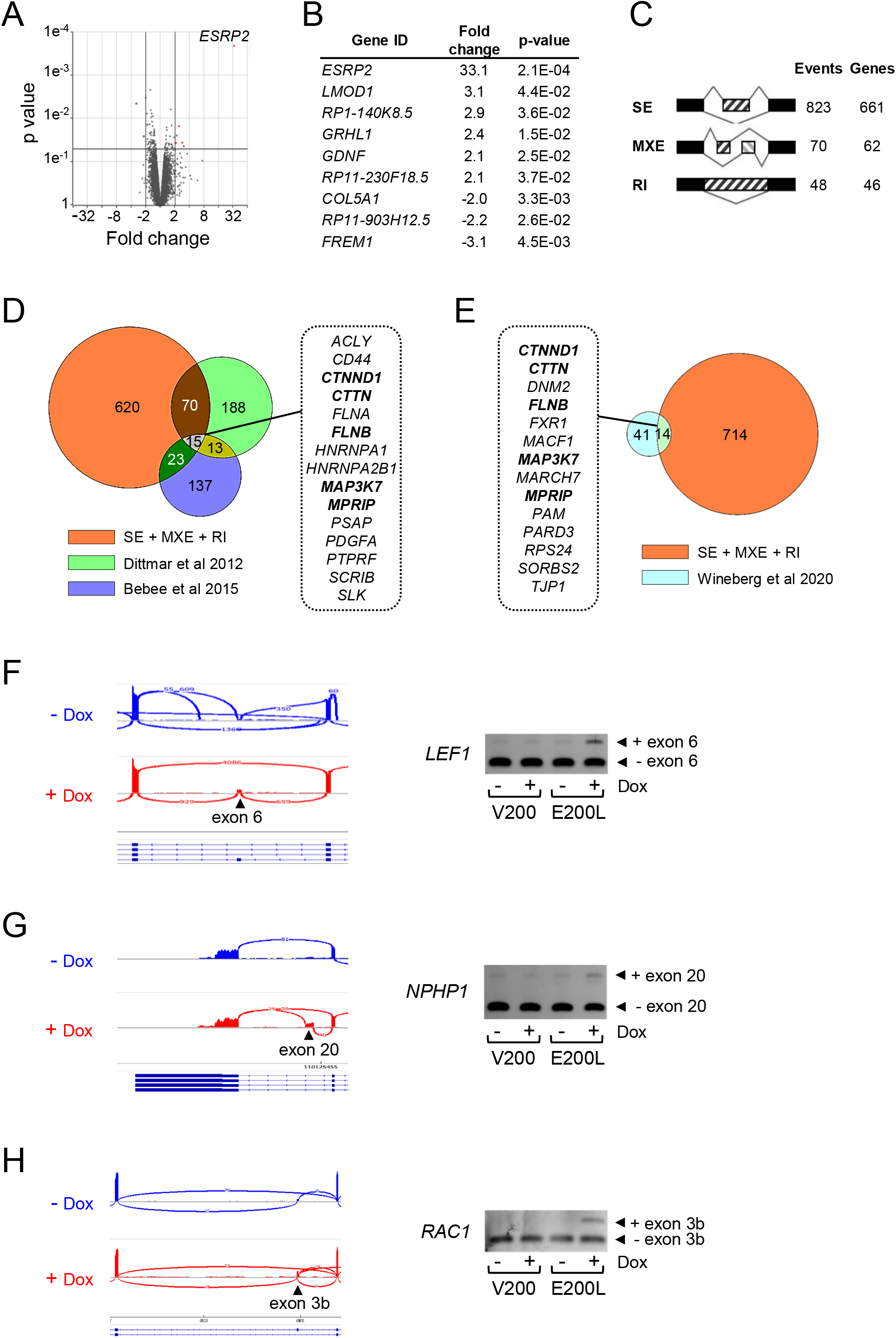
RNA-seq analysis. RNA-seq was performed on E200L cells with or without 96 hrs of Dox induction (to induce high-level ESRP2 expression). A: Volcano plot of p value versus fold induction of transcripts in ESRP2-expressing E200L cells compared to non-expressing cells. Genes induced >2 fold with p<0.05 are indicated in red, and *ESRP2* is labelled. B: List of genes shown in A that were induced >2 fold with p<0.05. C: Number of altered splicing events and affected genes induced by ESRP2 expression. SE; skipped exons, MXE; mutually exclusive exons; RI, retained introns. See supplementary tables S4, S5 and S6 for full details. D: Venn diagram comparing unique genes identified in this study (SE+MXE+RI) with two other RNA-seq analyses of ESRP-induced splicing changes (58, 59). E: Venn diagram comparing unique genes identified in this study (SE+MXE+RI) with a RNA-seq analysis of MET-associated splicing changes in the developing kidney (60). F, G and H: Alternative splicing of novel targets *LEF1* (F), *NPHP1* (G) and *RAC1* (H). Left-hand panels: Sashimi plots of RNA-seq data from E500L cells uninduced (-Dox) or induced to express ESRP2 (+Dox). Right-hand panels: Agarose gels of RT-PCRs of amplicons spanning the alternatively spliced exons (see supplementary table S9 for primers), in V200 and E200L cells, either uninduced, or Dox-induced to produce high-level ESRP2 expression in E200L cells.

In contrast to the lack of altered gene transcription, ESRP2 induction was associated with over 900 altered splicing events involving over 700 genes, with significant changes (False Discovery Rate, FDR<0.05) in skipped exons (SE), mutually exclusive exons (MXE) and retained introns (RI) (figure 6C, supplementary tables S4, S5, S6). The genes involved in these alternative splicing events were particularly enriched for biological processes concerned with vesicular and intracellular transport (supplementary table S7). There have been several previous studies identifying ESRP target genes in different biological contexts. Compared to other reports identifying specific targets of ESRPs using RNA-seq (58, 59), we found several targets in common, including well-known genes such as *CD44, CTNND1, SCRIB* and *SLK*, as well as over 600 genes that had not been previously described as ESRP2 targets (figure 6D). An additional comparison with a recent study of MET-associated alternative splicing changes during kidney development (60) also revealed overlap with some of our target genes (figure 6E). Interestingly, the two lists of genes identified in figures 6D and E as overlapping our ESRP2 targets, included five genes in common (*CTNND1, CTTN, FLNB, MAO3K7, MPRIP;* shown in bold in figures 6D and E). This extensive overlap (33-36%) between ESRP2 targets and genes showing splicing changes during kidney MET, emphasises the importance of ESRP-regulated alternative splicing in kidney development.

We first validated the putative target genes identified by RNA-seq, by using specific RT-PCR assays across alternatively spliced exons, to examine whether exon inclusion differed upon ESRP2 induction in the E200L cell line. We successfully validated several previously identified targets; *CD44, ENAH, FGFR2, SCRIB* and *SLK* (supplementary figure S12), as well as the novel targets *LEF1, NPHP1* and *RAC1* (figure 6F, G, H). However, some putative target genes showed no altered splicing after ESRP2 induction (supplementary figure S13). Overall, using the E200L cell line system, out of a total of 12 splicing events tested, 8 (67%) were validated. Seven of these events were novel, of which 3 (43%) were validated (supplementary table S8).

To investigate the possible role of ESRP2 target genes in WT pathogenesis, we examined alternative splicing of 12 genes (seven novel and five previously described), using the RT-PCR assays in fetal kidney, normal kidney and Wilms tumour (figure 7 and supplementary figure S14). Five genes showed significantly changes in the degree of alternative splicing between normal tissues and WT (figure 7), including one novel gene, *LEF1.* Seven genes did not show significant changes (supplementary figure S14), of which six were novel genes. Overall, 42% of the genes tested (five out of 12) demonstrated alternative splicing changes in WT, of which four were previously described targets (supplementary table S8).

**Figure 7:**
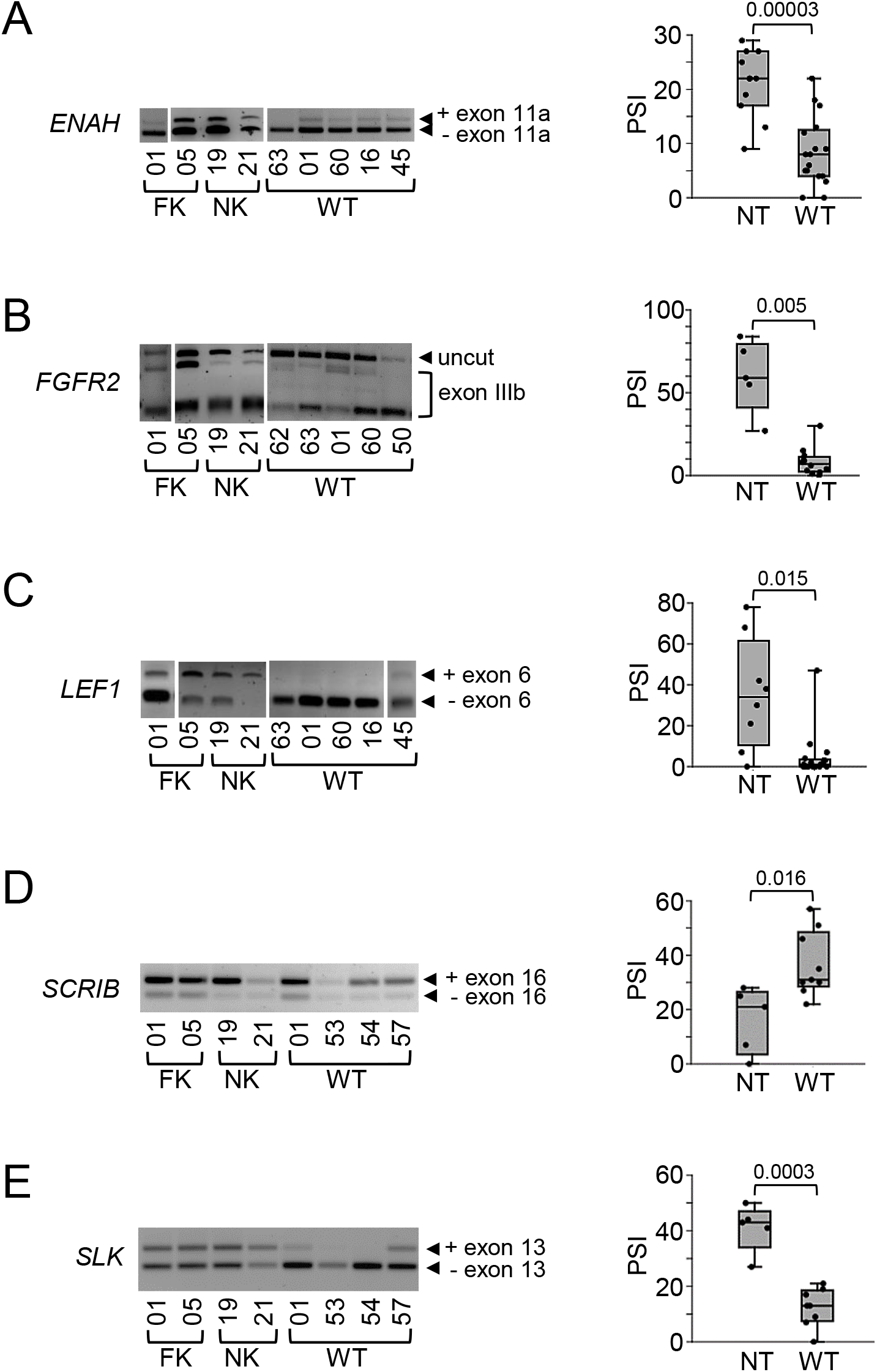
Alternative splicing of ESRP2 target genes in Wilms tumour. A to E: Left-hand panels: Representative agarose gels of RT-PCRs of amplicons spanning the alternatively spliced exons (see supplementary table S9 for primers), from FK, NK and WT. *FGFR2* exon IIIb was detected by restriction digest with *AvaI* (40). Right-hand panels: Dot-boxplots showing percent splice inclusion (PSI) in NT and WT. p values from t test. *ENAH* (A), NT n=11 (4 FK and 7 NK), WT n=17; *FGFR2* (B), NT n=5 (3 FK and 2 NK), WT n=12; *LEF1* (C), NT n=8 (5 FK and 3 NK), WT n=17; *SCRIB* (D), NT n=5 (2 FK and 3 NK), WT n=9; *SLK* (E), NT n=5 (2 FK and 3 NK), WT n=8.

## DISCUSSION

In this paper we have used genome-wide DNA methylation analysis to identify over 200 WT-associated epigenetic alterations. GO analysis showed that the hypermethylated genes were enriched in genes involved in developmental processes (figure 1B). Filtering of our gene list to remove those likely to be commonly epigenetically repressed in cancer, and to identify genes vital for early kidney development, highlighted four genes, one of which, *ESRP2*, was shown to be frequently hypermethylated in WT in an initial small-scale validation (figure 1C). Further investigations in two large independent WT cohorts demonstrated that *ESRP2* was hypermethylated in over 70% of WTs, which was confirmed using a publicly available dataset (figure 2A to C). Hypermethylation of *ESRP2* was associated with down-regulation of *ESRP2* expression, demonstrated in the two independent WT cohorts and another publicly available dataset (figure 3A to D). A mechanistic link between *ESRP2* DNA methylation and expression was further strengthened by the pharmacological reactivation of *ESRP2* by the methyltransferase inhibitor AZA in two hypermethylated WT cell lines (figure 3E). This is the first demonstration of loss of *ESRP2* expression caused by DNA methylation in Wilms tumour, and implicates RNA splicing alterations as an important pathogenic factor in WT development.

Investigation of *ESRP2* DNA methylation in two matched sets of NK, nephrogenic rest (NR) and WT, showed that *ESRP2* hypermethylation was at similar levels in NR and WT (figure 2D). This was confirmed in a publicly available dataset containing a much larger number of NRs (figure 2E). We also confirmed previous observations that *RASSF1A,* an epigenetically repressed tumour suppressor gene, is only hypermethylated in WTs, not in NRs. These results suggest that inactivation of *ESRP2* by DNA methylation occurs prior to NR formation, at an early stage in kidney development. This contrasts with *RASSF1A,* which is only hypermethylated in WTs, suggesting that it is epigenetically inactivated after NR formation. Mechanistically, the timing of *ESRP2* DNA methylation leads to a model in which ESRP2 is involved in early events in kidney development that are essential for the normal epithelial differentiation of the metanephric blastema into nephrons (figure 8B). Therefore, loss of ESRP2 expression due to DNA hypermethylation causes a differentiation block. This block initiates the persistence of undifferentiated renal precursors, which may then grow into a NR, that can then undergo further genetic and epigenetic defects to produce a WT (figure 8C). Support for this model comes from previous studies that have shown that the *Esrp* paralogs are expressed in the developing kidney (61) and that knockout of *Esrp* genes in mice decreases kidney volume, due to a lack of nephrons (62).

**Figure 8:**
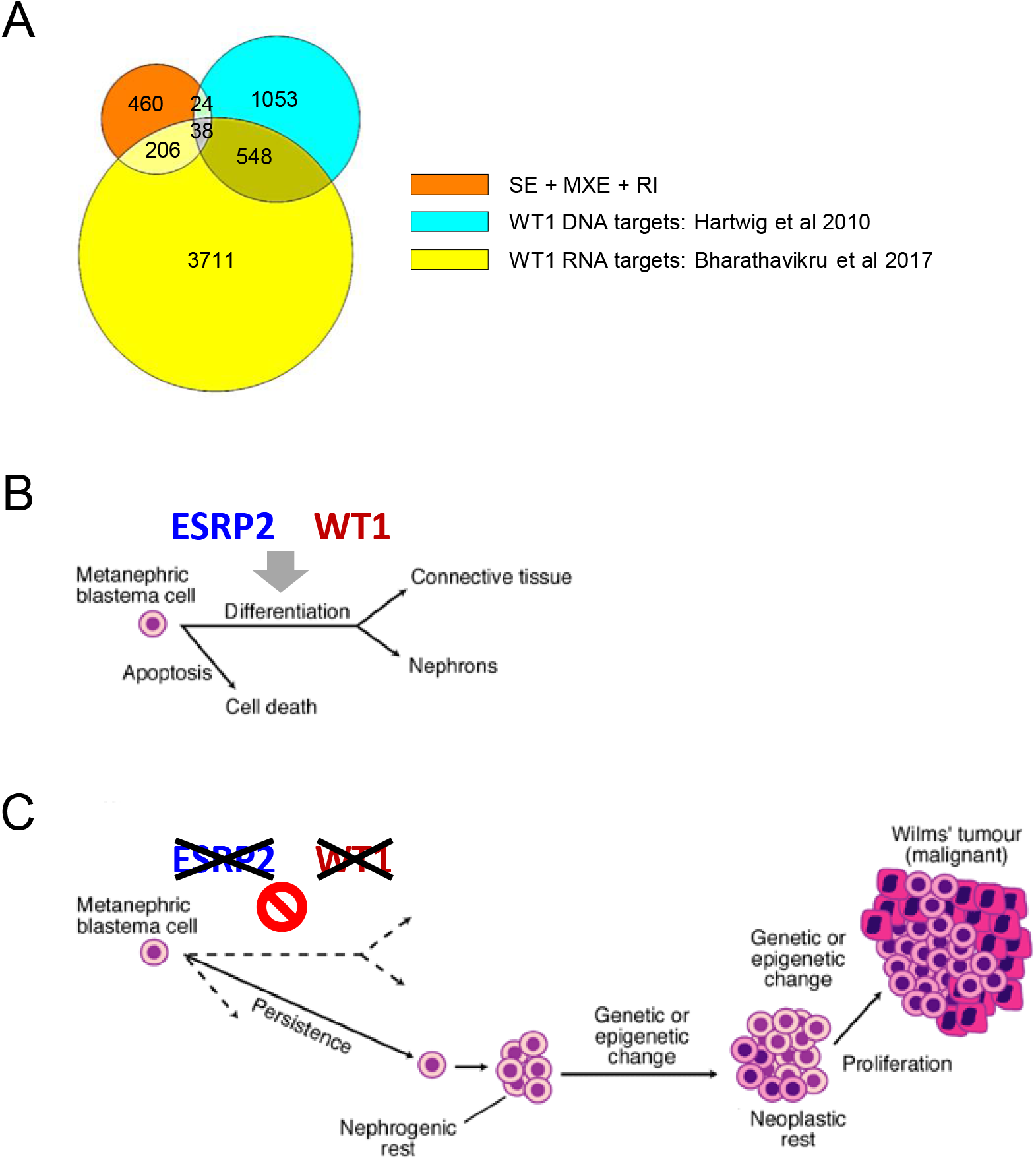
ESRP2 action in Wilms tumour. A: Venn diagram comparing the 728 unique genes identified in this study (SE+MXE+RI; figure 7) with 1663 WT1 DNA-binding targets identified by chromatin immunoprecipitation in developing kidney (82) and 4503 WT1 RNA-binding targets (protein-coding genes) identified by RNA immunoprecipitation in M15 mesonephric cells (83). B: ESRP2 and WT1 are required for epithelial differentiation, to form nephrons, during kidney development. C: Loss of ESRP2 function by hypermethylation or loss of WT1 function by mutation, inhibits normal differentiation and therefore promotes persistence of undifferentiated blastema, leading to nephrogenic rest formation and eventual progression to Wilms tumour. B and C adapted from figure 2 in reference (1).

The frequent epigenetic inactivation of *ESRP2* in early WT development, means that most WTs acquire this epigenetic lesion as a premalignant event, which probably explains why we found no association with clinical features of the WTs studied (supplementary figures S5 and S6). It also explains why we found no association between *ESRP2* methylation and LOH at 16q (supplementary figure S5C), where the *ESRP2* gene is located, because we have previously demonstrated that 16q LOH is a relatively late event in WT pathogenesis, occurring after NR formation (21) i.e. after *ESPR2* hypermethylation has occurred.

Interestingly, when we clustered WTs by DNA methylation at four loci, including *ESRP2,* we found no clinical associations, but we did find that all tumours with *WT1* mutations were in the same cluster, in which the WTs had relatively low levels of *ESRP2* DNA methylation (figure 2F). A recent comprehensive characterisation of genetic and epigenetic defects in a large series of WTs, also found that all *WT1*-mutant tumours were in a distinct group, based on DNA genome-wide DNA methylation analysis (14). We have shown here that *ESRP2* hypermethylation is an early event, being found in NRs (figure 2D, E), like *WT1* mutations (63, 64). These results therefore suggest that *WT1* mutation and *ESRP2* hypermethylation could be alternate early events in WT development, each of which can contribute to WT pathogenesis by inhibiting MET (figure 8B, C).

We have also found that *ESRP2* hypermethylation occurs in other childhood renal tumours (supplementary figure S7). Premalignant lesions analogous to NRs are unavailable for these tumours, so we have no way of knowing if *ESRP2* hypermethylation is an early or late event in the pathogenesis of these cancers. The different histology of these childhood renal tumours (65), together with more recent evidence on their cellular origins (66, 67), suggest that whilst *ESRP2* plays a role in their development, it is unlikely that their pathogenesis involves an MET block identical to that seen in WT.

We have shown that *ESRP1*, though an *ESRP2* paralogue, is not hypermethylated in WTs (figure 2B, C), and its expression is not repressed in WTs (figure 3C, D), suggesting that *ESRP1* and *ESRP2* may have different biological functions and are regulated differently in some instances. Similarly, a recent paper (68) has reported that only *ESRP2* and not *ESRP1* is regulated by androgens, with important implications in prostate cancer progression.

Splicing alterations are frequent in human cancers (69), including changes induced by *ESRP* genes, such as in breast cancer (54, 70), prostate cancer (68, 71), renal cell carcinoma (72) and colorectal cancer (73). In most cases, *ESRP* genes appear to have tumour suppressive properties, presumably because loss of *ESRP* expression permits EMT (27, 39), which is important in many stages of carcinogenesis, especially the acquisition of malignant characteristics i.e. invasion and metastasis. (74). This contrasts with the results reported in this paper, where loss of *ESRP2* function appears to inhibit MET at a very early, premalignant stage of tumour development.

Most studies on *ESRP* genes in human cancer demonstrate expression changes but without any reported underlying genetic or epigenetic defects in the *ESRP* genes themselves (54, 68, 70, 72, 73). However, there have been reports of genetic defects in *ESRP* genes in human cancers, specifically, microsatellite indels (75) or duplications (71) of *ESRP1.* In addition, there have been reports of DNA methylation changes of *ESRP1* in prostate cancer (76) and of *ESRP2* in breast cancer (77). Thus, our results add to a growing body of evidence that *ESRP* genes can be either genetically or epigenetically deregulated in a wide range of human cancers.

Functionally, our studies with a WT cell line carrying an inducible *ESRP2* construct, suggest that the main biological effect of ESRP2 is to regulate cell proliferation by slowing cell division (figure 4H to K and supplementary figure S8). Whilst we observed some actin cytoskeleton rearrangement (figure 4 D to G), we did not observe significant expression changes in classical epithelial marker genes (figure 6A, B), nor any changes in cell motility or invasion (supplementary figure S9), as have been reported in studies where ESRP expression has been modulated in adult human cancer cell lines (39, 54, 55). Coupled with our xenograft experiments that identify *ESRP2* as a *bona fide* tumour suppressor gene in a WT line cell (figure 5), these results suggest that the tumour suppressor activity of *ESRP2* in WT occurs mainly by altering cell growth properties, rather than by affecting cellular differentiation.

Mechanistically, our RNA-seq results demonstrate that ESRP2 modulates the splicing of a diverse range of genes, including both well-established and novel targets (figure 6 and supplementary tables S4 to S6). We showed that a subset of these genes demonstrate reduced expression of their epithelial splice forms in WT (figure 7), consistent with the DNA-methylation induced down-regulation of *ESRP2* that we have observed in the majority of WTs (figures 2 and 3). Interestingly, *LEF1, a* novel ESRP2 target gene that shows deregulated splicing in WTs (figure 6F and 7C), is a component of the Wnt signalling pathway (78) and is important for the differentiation of nephron progenitor cells during kidney development (79). Disruption of the Wnt pathway has been identified as central to WT pathogenesis (3), with mutations observed in the Wnt pathway genes *CTNNB1* (6, 7) and *WTX* (*AMER1*) (8) and *Wt1* shown to be essential in the control of *Wnt4* during normal nephrogenesis (80). Thus, epigenetic inactivation of *ESRP2*, as described here, could elicit an early defect in the Wnt pathway via its modulation of *LEF1* splicing (81), with similar effects to the loss of *WT1* early in WT pathogenesis.

Interestingly, of the 728 genes that we identified as having their splicing modulated by ESRP2 (figure 6), only 62 (9%) are WT1 DNA-binding targets (82), whereas 244 (34%) are WT1 RNA-binding targets (83) (figure 8A). The WT1 RNA-binding targets include all five of the ESRP2-regulated genes that we found in common between our results and two other RNA-seq studies (figure 6D, E). This suggest that post-transcriptional regulation of gene expression, either by ESRP2 or WT1, may be alternative mechanisms for regulating important renal developmental genes. This could explain the different DNA methylation profiles that we observed in WTs with and without *WT1* mutations (figure 2F), suggesting that epigenetic inactivation of *ESRP2* and mutation of *WT1* may be alternate events that control an early stage of WT development (figure 8B, C). These results, together with genetic evidence showing defects in miRNA-processing genes in WT (9–13), reinforce the critical role that post-transcriptional gene regulation plays in WT pathogenesis.

## Abbreviations

Aza: 5-aza-2’-deoxycytidine
BCH: Bristol Children’s Hospital
CCSK: clear cell sarcoma of the kidney
CGI: CpG island
CTV: CellTrace Violet
DMEM: Dulbecco’s modified Eagle’s medium
Dox: doxycycline
ES: embryonic stem cells
FBS: fetal bovine serum
FK: fetal kidney
MCIP: methyl CpG immunoprecipitation
MET: mesenchymal-epithelial transition
NK: normal kidney
NR: nephrogenic rest
NT: normal tissue
PBS: phosphate buffered saline
PRC: polycomb repressive complex
QPCR: quantitative real-time PCR
RH: rhabdoid tumour
RMH: Royal Marsden Hospital
RNA-seq: RNA sequencing
STR: short tandem repeat
WT: Wilms tumour

## Acknowledgments

The authors thank: Professor Herman Yeger for the gift of the Wit49 cell line; Dr Madhu Kollareddy for the STR profile of Wit49; the Bristol Genetics Laboratory, Southmead Hospital, for help with pyrosequencing; Dr Paul Bishop for advice about immunofluorescence; Dr. Andrew Herman and Lorena Sueiro Ballesteros for the flow cytometry; and the University of Bristol Genomics Facility for the RNA sequencing work.

## Author contributions

D Legge carried out the RNA-seq, target validation and other experimental work, LL performed the animal experiments, WM carried out pyrosequencing, QPCR and derived the inducible cell line, D Lee performed the bioinformatic analysis of the RNA-seq data, MS performed the MCIP DNA methylation analysis, AZ, LP and WYC performed DNA methylation and expression analyses on cell lines and tumours, YA and JB validated some putative ESRP2 targets, RW and KPJ provided tumour samples and clinical data, KTAM help devise the methylation analysis strategy, SO planned experiments, especially the animal work and KWB wrote the paper, planned work and carried out some experimental work. All authors viewed the manuscript and were given the opportunity to comment on it.

