## Supplementary figures for "The epithelial splicing regulator *ESRP2* is epigenetically repressed by DNA hypermethylation in Wilms tumour and acts as a tumour suppressor"

**Legge et al**

### Supplementary data: Figures S1 to S24

| | |
|---|---|
| **S1** | Cell line STR profiles |
| **S2** | *ESRP2* inducible expression construct |
| **S3** | *ESRP2* methylation detected by MCIP |
| **S4** | *ESRP2* methylation and RNA expression in Wilms tumours from two cohorts |
| **S5** | *ESRP2* DNA methylation in Wilms tumours of different stages and 16q LOH status |
| **S6** | Overall survival in Wilms tumour patients with different levels of *ESRP2* methylation |
| **S7** | *ESRP2* methylation in non-Wilms childhood renal tumours |
| **S8** | Growth of Wit49 transfected cells |
| **S9** | Motility assays of Wit49 transfected cells |
| **S10** | Mouse tumorigenicity primary data |
| **S11** | *GRHL1* RNA expression |
| **S12** | Successfully validated putative ESRP2 target genes |
| **S13** | Unsuccessfully validated putative ESRP2 target genes |
| **S14** | Alternative splicing of putative ESRP2 target genes in normal kidney and Wilms tumour |
| **S15** | Uncropped gels for figure 1 |
| **S16** | Uncropped blots and gel for figure 4 |
| **S17** | Uncropped blots for figure 5 |
| **S18** | Uncropped gels for figure 6 |
| **S19** | Uncropped gels for figure 7 |
| **S20** | Uncropped gels for figure S1B |
| **S21** | Uncropped gels and blots for figure S2B and S2C |
| **S22** | Uncropped gels for figure S12 |
| **S23** | Uncropped gels for figure S13 |
| **S24** | Uncropped gels for figure S14 |

A

| Locus | Wit49 Yeger | Wit49 Bristol |
|---|---|---|
| AM | X | X, X |
| D3S1358 | 17 | 17, 17 |
| D1S1656 | | 12, 16 |
| D6S1043 | | 11, 12 |
| D13S317 | 11, 12 | 11, 12 |
| Penta E | 13 | 13, 13 |
| D16S539 | 12 | 12, 12 |
| D18S51 | 15 | 15, 15 |
| D2S1338 | | 23, 26 |
| CSF1PO | 11, 12 | 11, 12 |
| Penta D | 11, 12 | 11, 12 |
| TH01 | 9.3 | 9.3, 9.3 |
| vWA | 16, 18 | 16, 18 |
| D21S11 | 29, 32.2 | 29, 32.2 |
| D7S820 | 8, 11 | 8, 11 |
| D5S818 | 12, 13 | 12, 13 |
| TPOX | 8 | 8, 8 |
| D8S1179 | 10 | 10, 10 |
| D12S391 | | 20, 23 |
| D19S433 | | 14, 14 |
| FGA | 25 | 25, 25 |

B

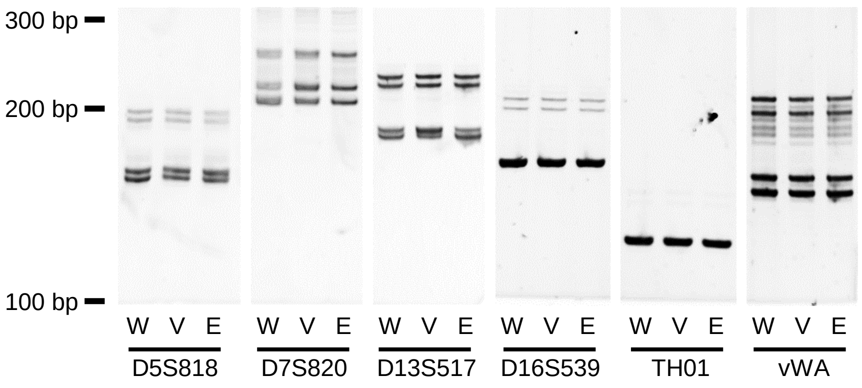


**Figure S1: Cell line STR profiles**
**A:** STR profiles for the original Wit49 cell line (personal communication Prof. H Yeger) and Wit49 cells (carried out by Eurofins; https://www.eurofinsgenomics.eu/) used in our laboratory (Bristol) for this paper.
**B:** Six STR markers amplified by PCR (supplementary table S9) and run on 7% non-denaturing acrylamide gels. Original Wit49 cells (W), and the V200 (V) and E200L (E) derivatives used in this paper all show identical profiles, with the STR amplicons having the predicted sizes from the previous STR profiling listed in A.

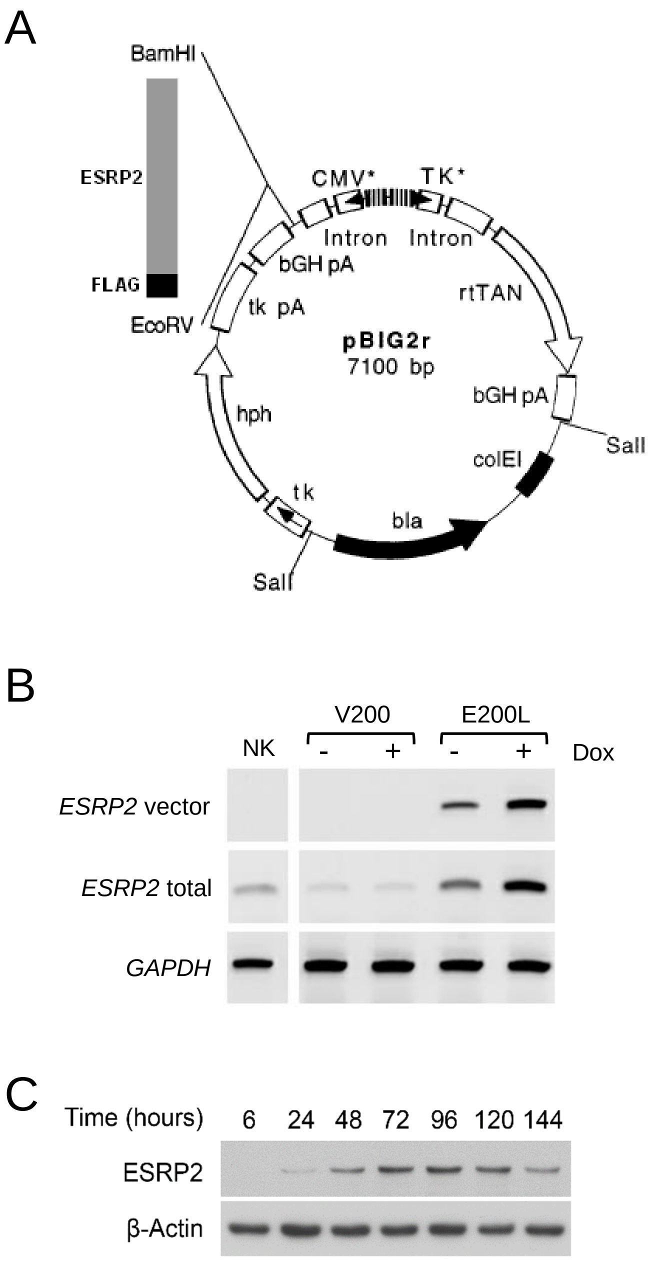


**Figure S2: *ESRP2* inducible expression construct**

**A:** *ESRP2* cDNA was amplified by PCR from IMAGE clone 4810948, using a forward primer containing a BamHI site and a reverse primer containing an EcoRV site plus a FLAG tag (supplementary table S9). This insert was then ligated into BamHI/EcoRV-digested pBIG2r (Strathdee, C. A. et al (1999) Gene 229(1-2): 21-29).

**B:** RT-PCR of cDNA from normal kidney (NK), control vector-transfected Wit49 cells (V200) and *ESRP2*-transfected Wit49 cells (E200L), amplified with primers for *ESRP2* expressed from the vector (ESRP2 RTF2 and BGHR), for total *ESRP2* (ESRP2 RTF2 and ESRP2 RTR2) and for *GAPDH* (GAPDH F and GAPDH R). Primer BGHR is located in the bGH pA region in the pBIG2R vector downstream of the insert (see A above) and the *ESRP2* primers are in the cDNA (supplementary table S9).

**C:** Western blot showing time course of ESRP2 protein expression induced in cell line E200L with 2 µg/ml doxycycline.

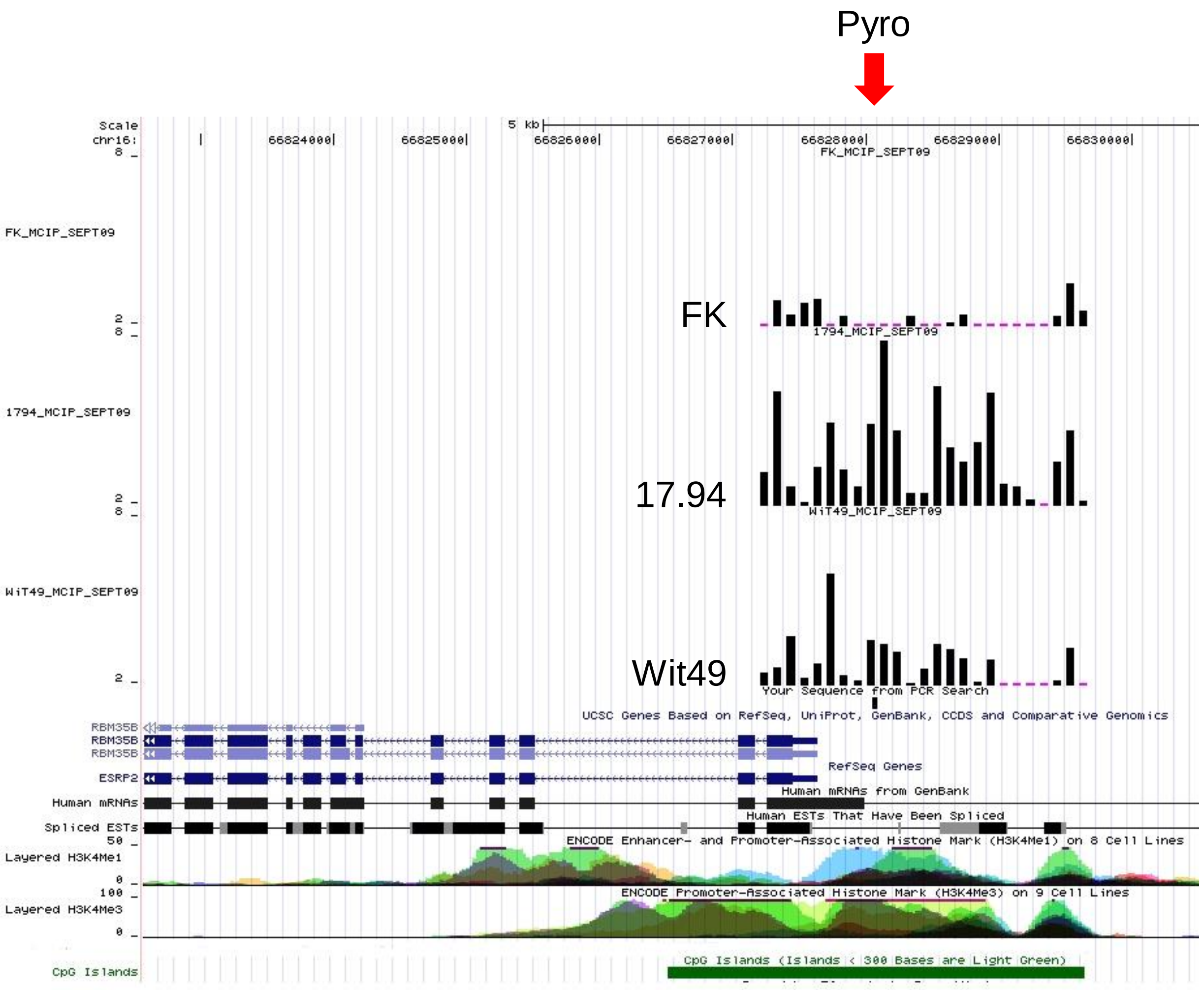


**Figure S3: *ESRP2* methylation detected by MCIP**
Black bars show the probe ratios derived from MCIP for fetal kidney (FK) and the two WT cell lines, 17.94 and Wit49, positioned on the *ESRP2* gene, showing the transcripts (*ESRP2* / *RBM35B*), H3K4Me1 and Me3 marks (ENCODE) and CpG islands (http://genome.ucsc.edu). The position of the pyrosequencing assay is indicated by the red arrow shown at the top.

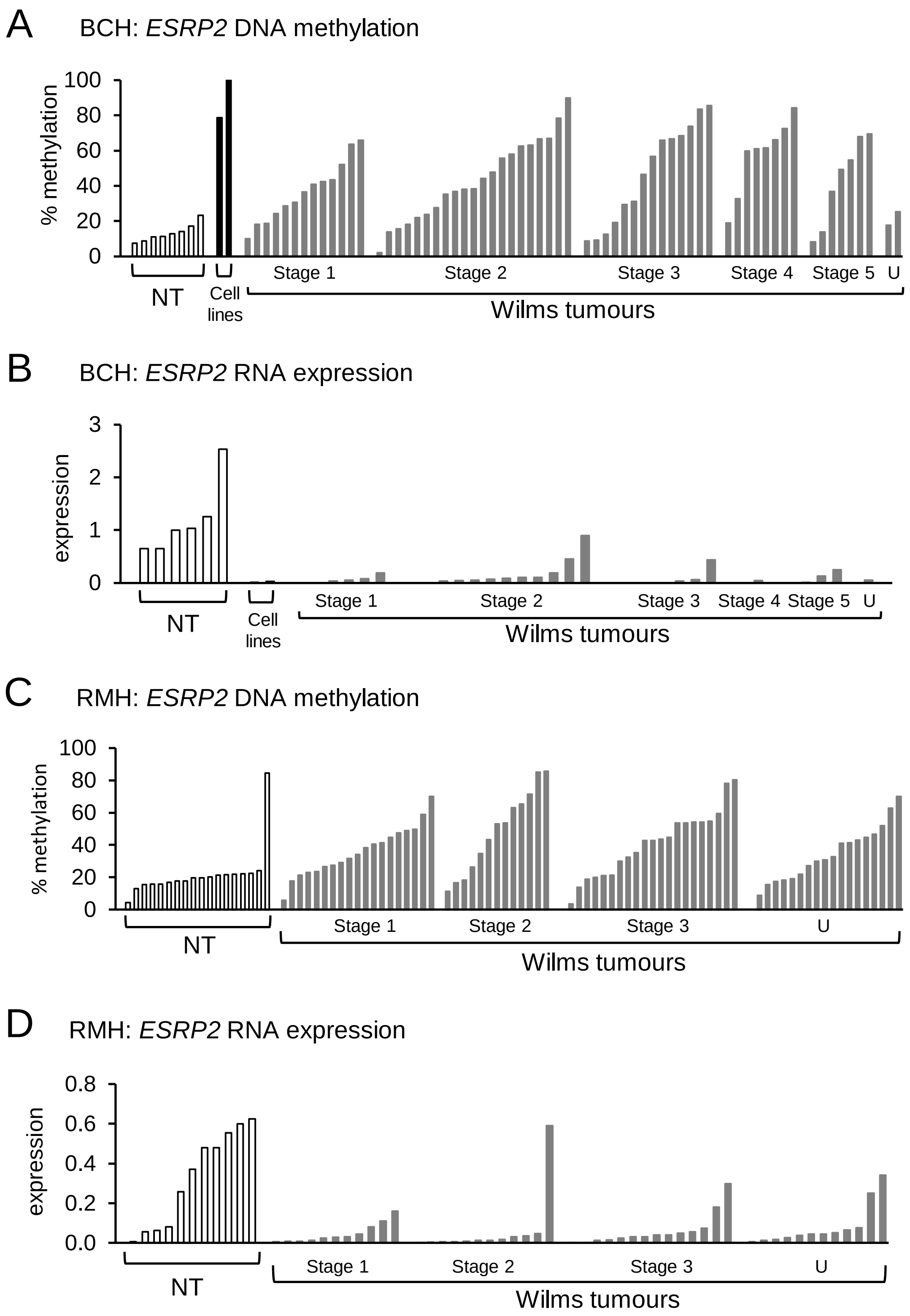


**Figure S4: *ESRP2* methylation and RNA expression in Wilms tumours from two cohorts**

**A:** *ESRP2* DNA methylation versus stage (BCH data; pyrosequencing).
**B:** *ESRP2* RNA expression versus stage (BCH data; QPCR).
**C:** *ESRP2* DNA methylation versus stage (RMH data; pyrosequencing).
**D:** *ESRP2* RNA expression versus stage (RMH data; QPCR).
NT, normal tissues; U, stage unknown.

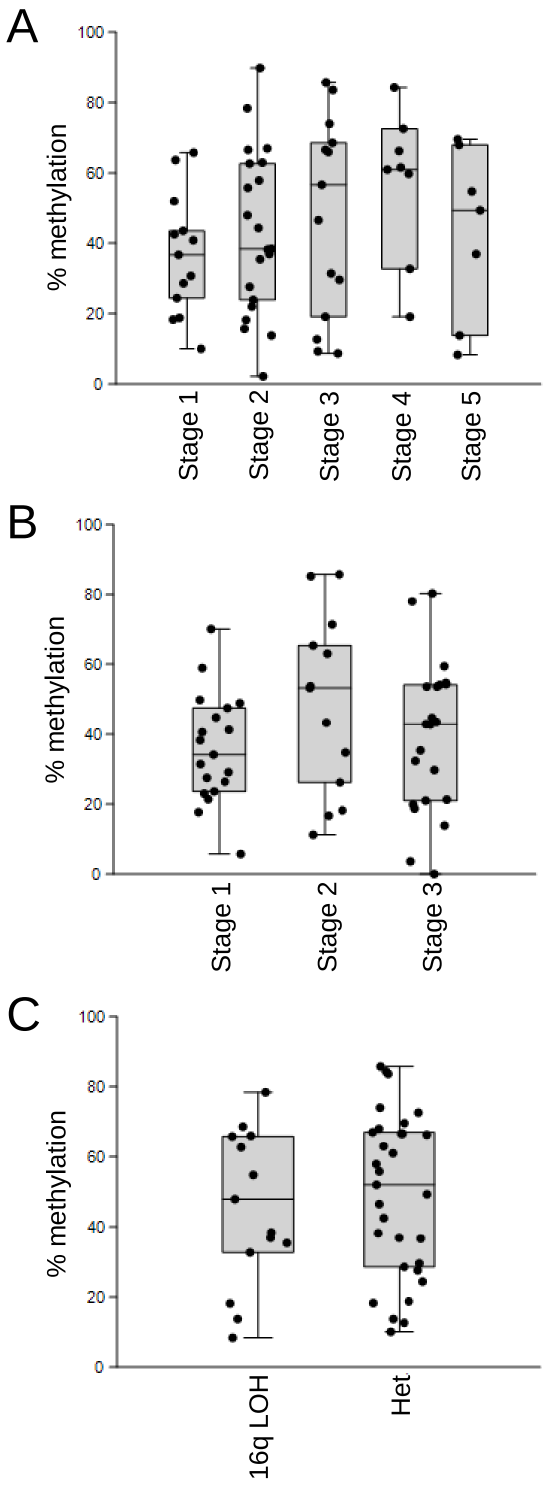


**Figure S5:** ***ESRP2*** **DNA methylation in Wilms tumours of different stages and 16q LOH status**

**A:** Dot-Boxplot of *ESRP2* DNA methylation versus stage (BCH data; for stages 1, 2, 3, 4, 5; n = 16, 24, 16, 9, 8 respectively).

**B:** Dot-Boxplot of *ESRP2* DNA methylation versus stage (RMH data; for stages 1, 2, 3; n = 13, 7, 16 respectively).

**C:** Dot-Boxplot of *ESRP2* DNA methylation versus 16q loss of heterozygosity (BCH data; LOH, n = 8; Het, n = 22). LOH, loss of heterozygosity; Het, heterozygous tumours.

*ESRP2* DNA methylation was assayed by pyrosequencing.

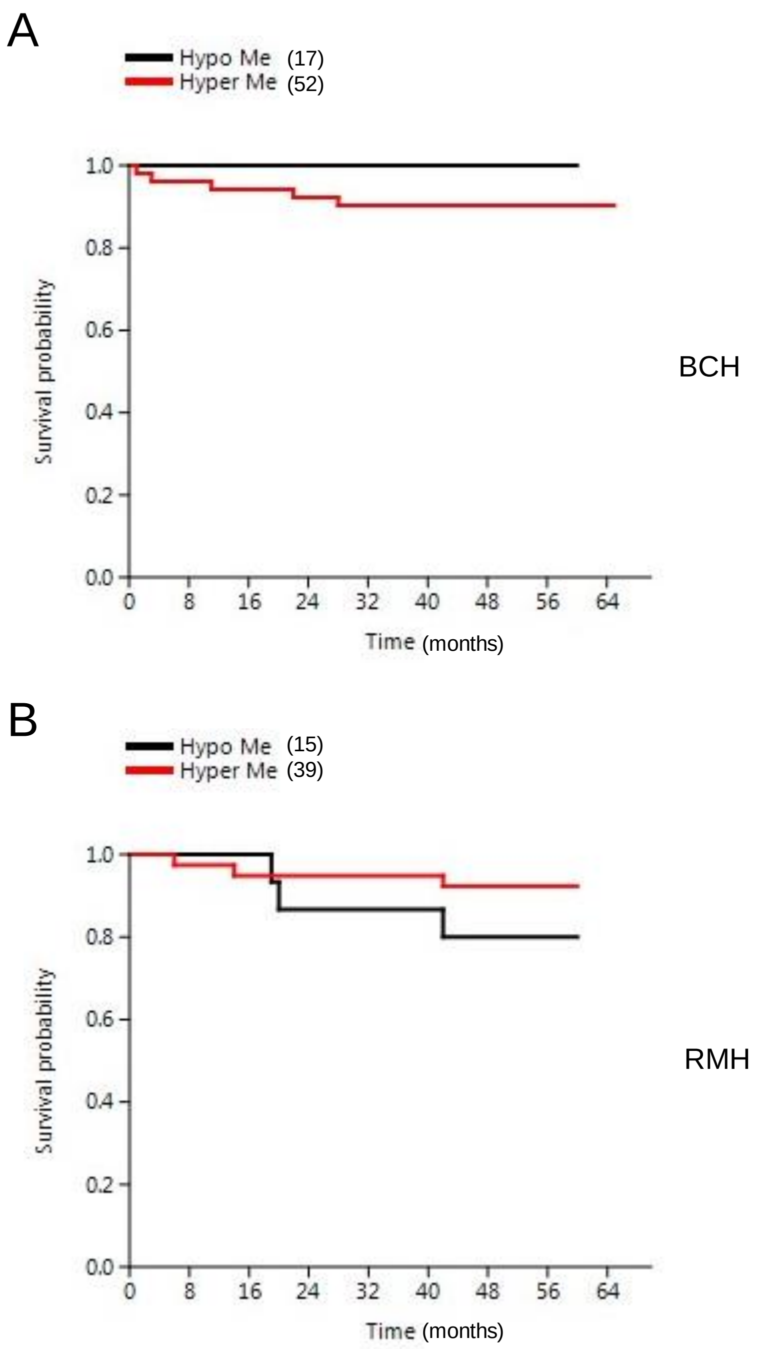


**Figure S6: Overall survival in Wilms tumour patients with different levels of *ESRP2* methylation**

**A:** Overall patient survival up to 5 years post diagnosis – BCH cohort. p = 0.200

**B:** Overall patient survival up to 5 years post diagnosis – RMH cohort. p = 0.212

Hypo Me, hypomethylated tumours (*ESRP2* DNA methylation <25%).

Hyper Me, hypermethylated tumours (*ESRP2* DNA methylation >25%).

*ESRP2* DNA methylation was assayed by pyrosequencing.

p values calculated using log-rank test.

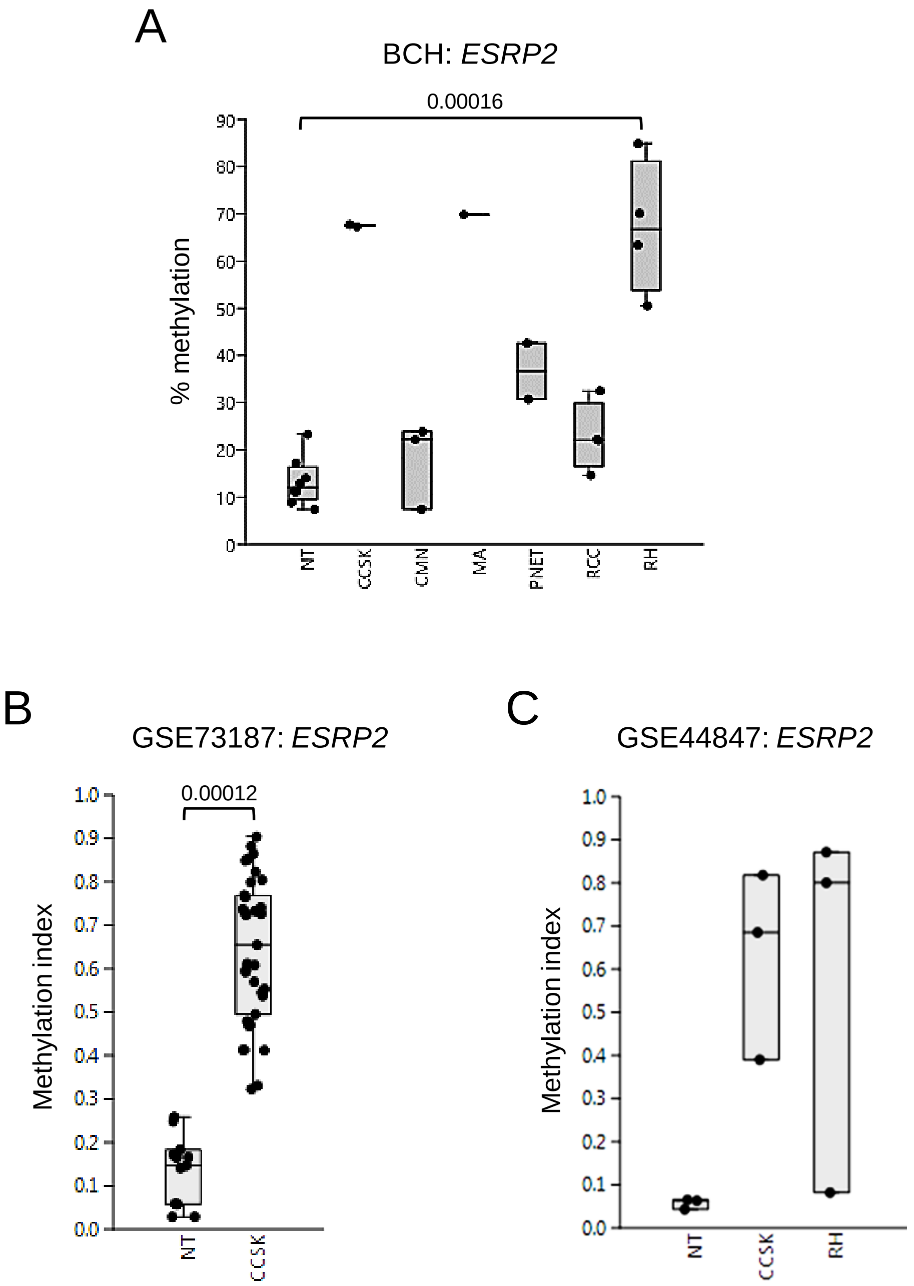


**Figure S7: *ESRP2* methylation in non-Wilms childhood renal tumours**

**A:** Dot-Boxplot of DNA methylation of *ESRP2* assayed by pyrosequencing in eight normal tissue (NT) samples (four fetal kidneys and four normal kidneys), two clear cell sarcomas (CCSK), three mesoblastic nephromas (CMN), one metanephric adenoma (MA), two primitive neuroectodermal tumours (PNET), four renal cell carcinomas (RCC) and four rhabdoid tumours (RH), from local samples (BCH).

**B:** Dot-Boxplot of methylation data from GSE73187 of 12 normal tissue (NT) samples (four fetal kidneys and eight normal kidneys) and 34 CCSKs.

**C:** Dot-Boxplot of methylation data from GSE44847 of three normal kidney (NT) samples, three CCSKs and three RHs.

In **B** and **C** samples were assayed by Illumina Human Methylation 27 BeadChip arrays. Data shown is from cg20264732, which is in the 5' CpG island of *ESRP2*. p value in A from Tukey's pairwise test, p value in B from t test.

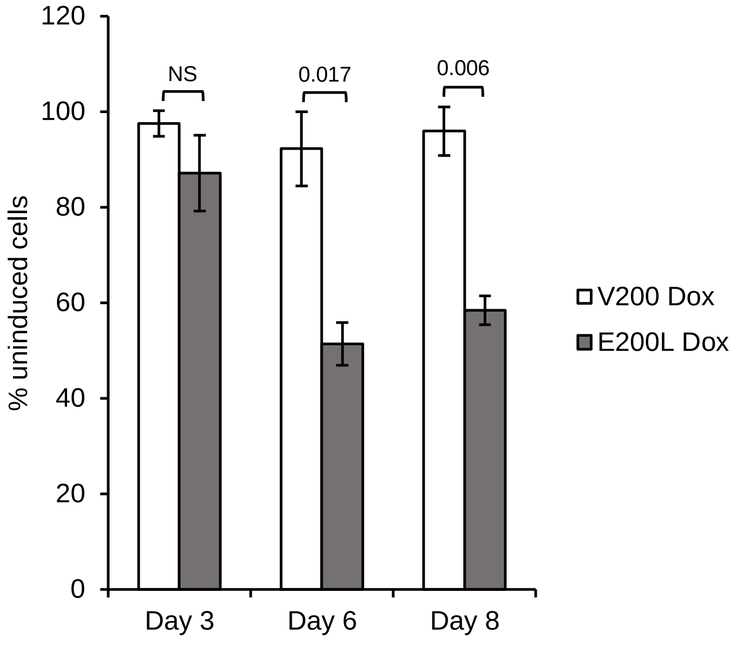


**Figure S8: Growth of Wit49 transfected cells**
Empty vector-transformed cells (V200) and *ESRP2*-transfected cells (E200L) were plated with and without doxycycline (2 µg/ml to induce *ESRP2* expression) and counted after 3, 6 and 8 days. Counts shown are the means ± SEM of three experiments, expressed as the percentage of the uninduced cells at each time point. Cell growth was significantly reduced at 6 days and at 8 days (t-test) in E200L cells compared to V200 control cells.

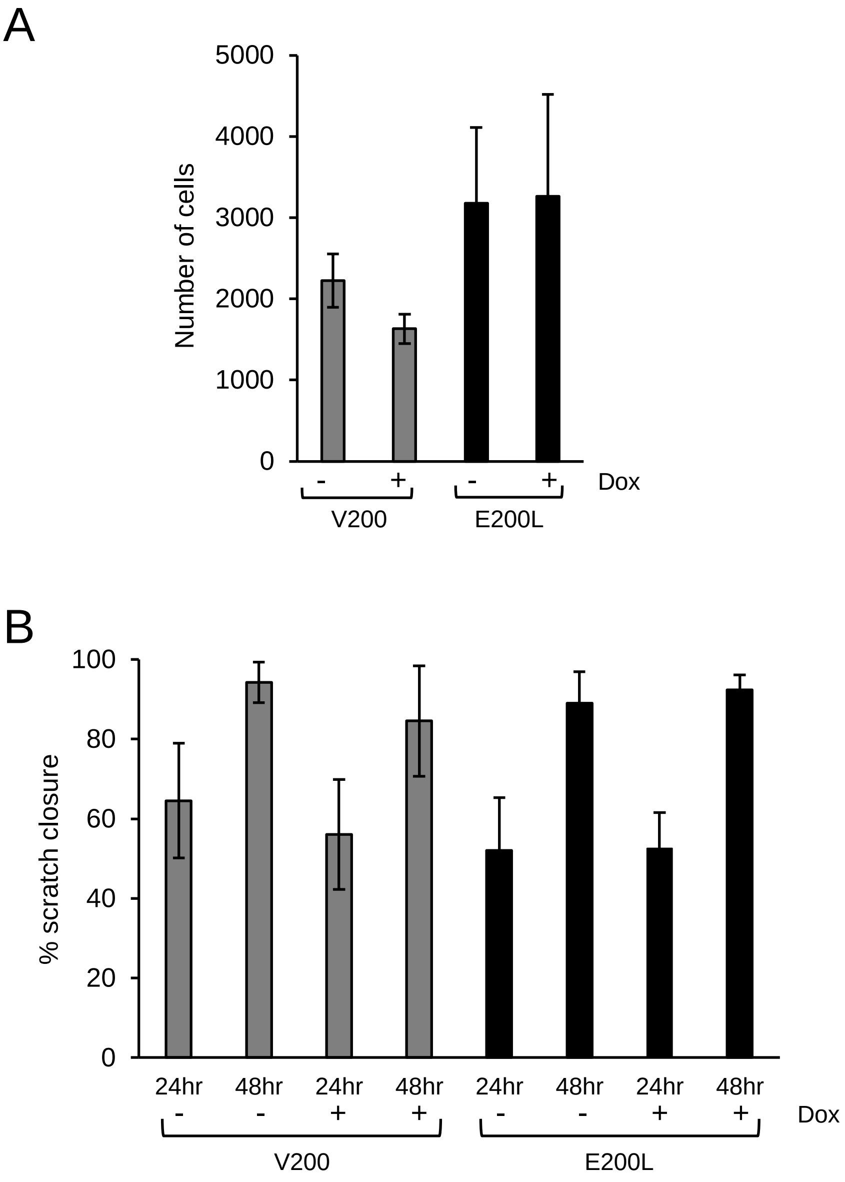


**Figure S9: Motility assays of Wit49 transfected cells**
**A:** Transwell assay; Cells were pre-treated for 4 days with 2 µg/ml Dox or control media, then seeded into transwell inserts in FBS-free DMEM and the inserts were put into wells filled with 10% FBS DMEM to produce a chemotactic gradient. Following 24 hours, inserts were washed, cells were fixed, stained and counted manually using light microscopy. Counts shown are the means ± SEM of three experiments.
**B:** Scratch assay; cells were seeded into 24 well plates and treated for 5 days with 2 µg/ml Dox or control media prior to a scratch being performed manually in the centre of each well. Wells were washed gently with PBS to remove dead cells from the scratch wound site. Control/Dox media was replaced and wells were analysed at 24 and 48 hours via widefield microscopy and Image J software to determine percentage wound closure. Counts shown are the means ± SEM of three experiments.

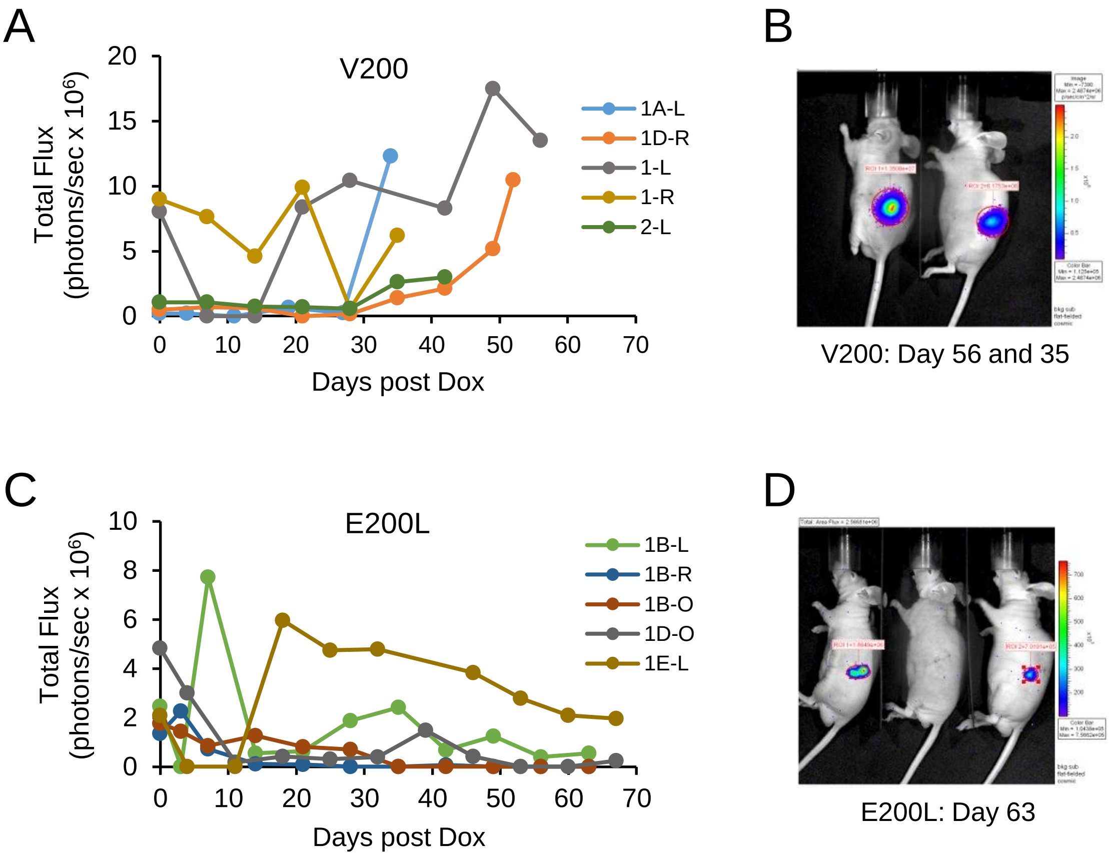


E

| Group | Mouse ID | Days after Doxy induction | Tumour width (mm) | Tumour length (mm) | Tumour volume (mm³) | Tumour Weight (mg) |
|---|---|---|---|---|---|---|
| V200 (Control) | 1A-L | 34 | 10 | 11 | 1155 | 345 |
| | 1D-R | 52 | 9 | 10 | 855 | 274 |
| | 1-L | 56 | 8 | 10 | 720 | 155 |
| | 1-R | 35 | 5 | 8 | 260 | 127 |
| | 2-L | 42 | 2 | 2 | 8 | 7 |
| E200L (ESRP2) | 1B-L | 63 | 3 | 4 | 42 | 35 |
| | 1B-R | 63 | 2 | 3 | 15 | 8 |
| | 1D-O (two tumours) | 67 | 1 | 2 | 3 | 5 |
| | | | 2 | 3 | 15 | 9 |
| | 1E-L | 67 | 9 | 10 | 855 | 257 |

**Figure S10: Mouse tumorigenicity primary data**
**A** and **C**: Time course of tumour growth as assayed by *in vivo* bioluminescence in V200 and E200L xenografts. Plots show tumour signals days after Dox induction (i.e. first Dox injection = day zero). **B** and **D**: Examples of *in vivo* bioluminescence imaging of nude mice carrying V200 and E200L xenografts. **E**: Size and weight of excised tumours.

A

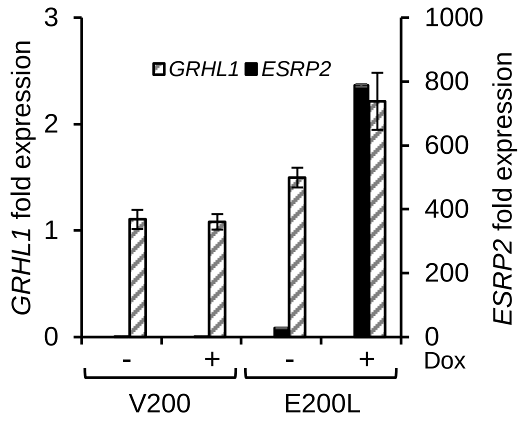


B

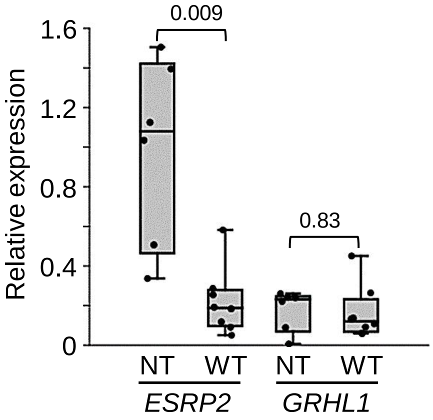


**Figure S11: *GRHL1* RNA expression**

**A:** QPCR of *ESRP2* and *GRHL1* RNA expression, normalized to endogenous levels of *TBP*, in Dox-induced and uninduced V200 and E200L cells. Expression shown as mean ± SEM of n=3, relative to uninduced V200 cells.

**B:** Dot-boxplot QPCR of *ESRP2* and *GRHL1* RNA expression, normalized to endogenous levels of *TBP*. NT n=6 (3 NK and 3 FK) and WT n=8, p values from t test.

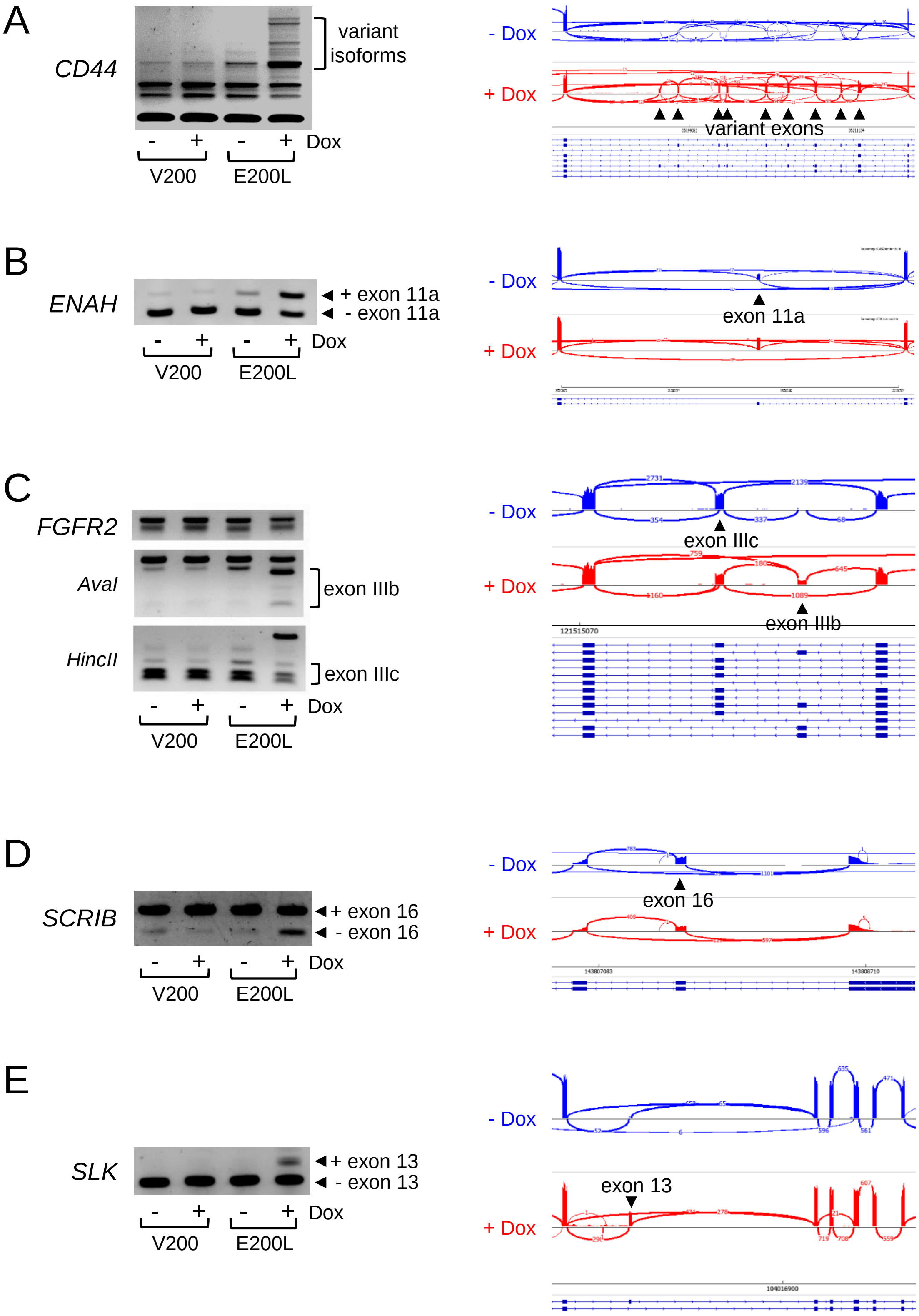


**Figure S12: Successfully validated putative ESRP2 target genes**
Left-hand panels: Target genes were amplified by RT-PCR for alternatively spliced exons (see supplementary table S9 for primers). *FGFR2* exons were detected by restriction enzyme cleavage (Warzecha et al (2009) Mol. Cell 33(5): 591-601) . Right-hand panels: Sashimi plots of RNA-seq data from E500L cells uninduced (-Dox) or induced to express ESRP2 (+Dox).

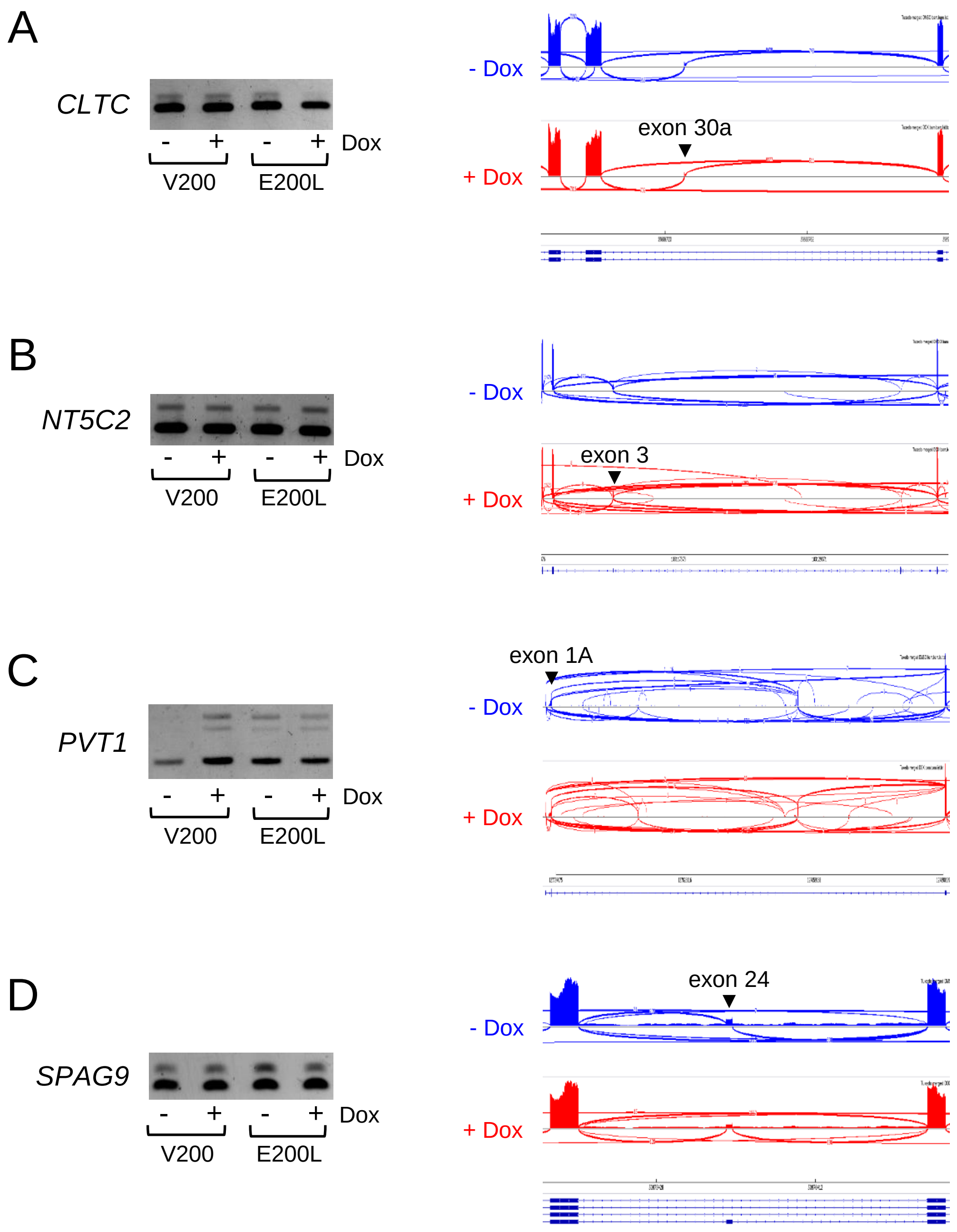


**Figure S13: Unsuccessfully validated putative ESRP2 target genes**
Left-hand panels: Target genes were amplified by RT-PCR for alternatively spliced exons (see supplementary table S9 for primers).
Right-hand panels: Sashimi plots of RNA-seq data from E500L cells uninduced (-Dox) or induced to express ESRP2 (+Dox).

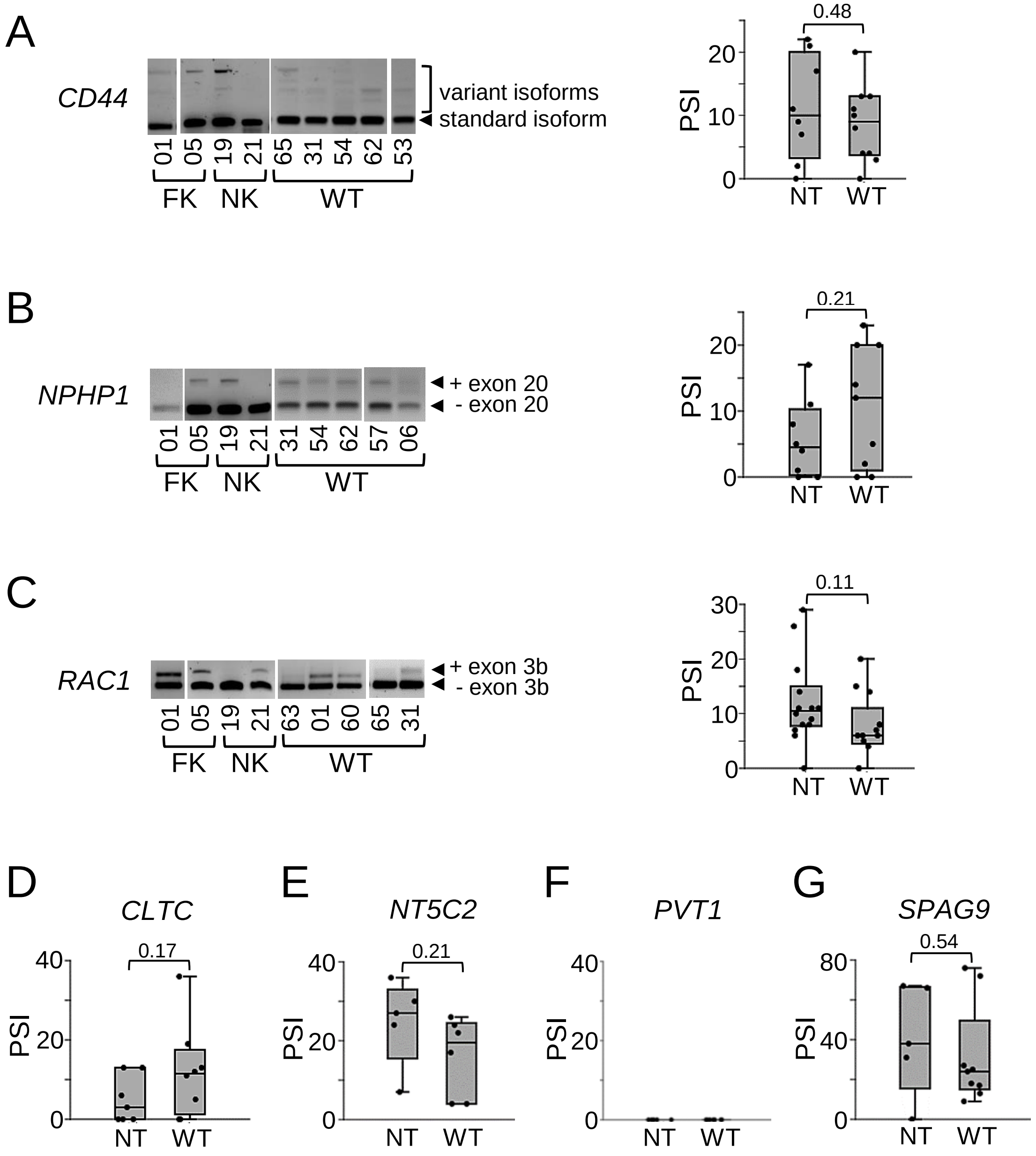


**Figure S14: Alternative splicing of putative ESRP2 target genes in normal kidney and Wilms tumour**

**A to C:** Left-hand panels: Target genes were amplified by RT-PCR for alternatively spliced exons (see supplementary table S9 for primers). Right-hand panels: Dot-Boxplots showing percent splice inclusion (PSI) in normal tissues (NT; NK and FK) and WT. p values from t test. **D to G:** Dot-boxplots showing percent splice inclusion (PSI) in normal tissues (NT; NK and FK) and WT. p values from t test.

D

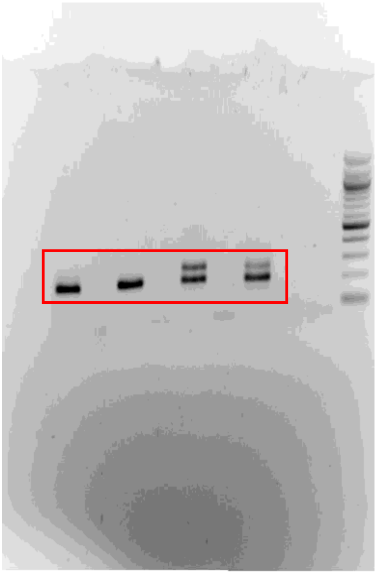

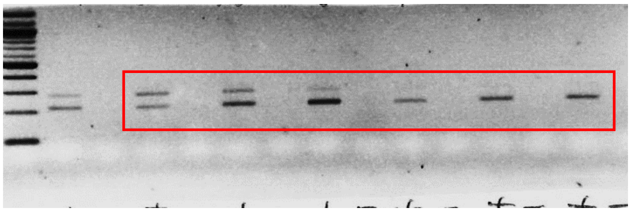

**Figure S15: Uncropped gels for figure 1**
Areas used in figure 1 are outlined in red.

B

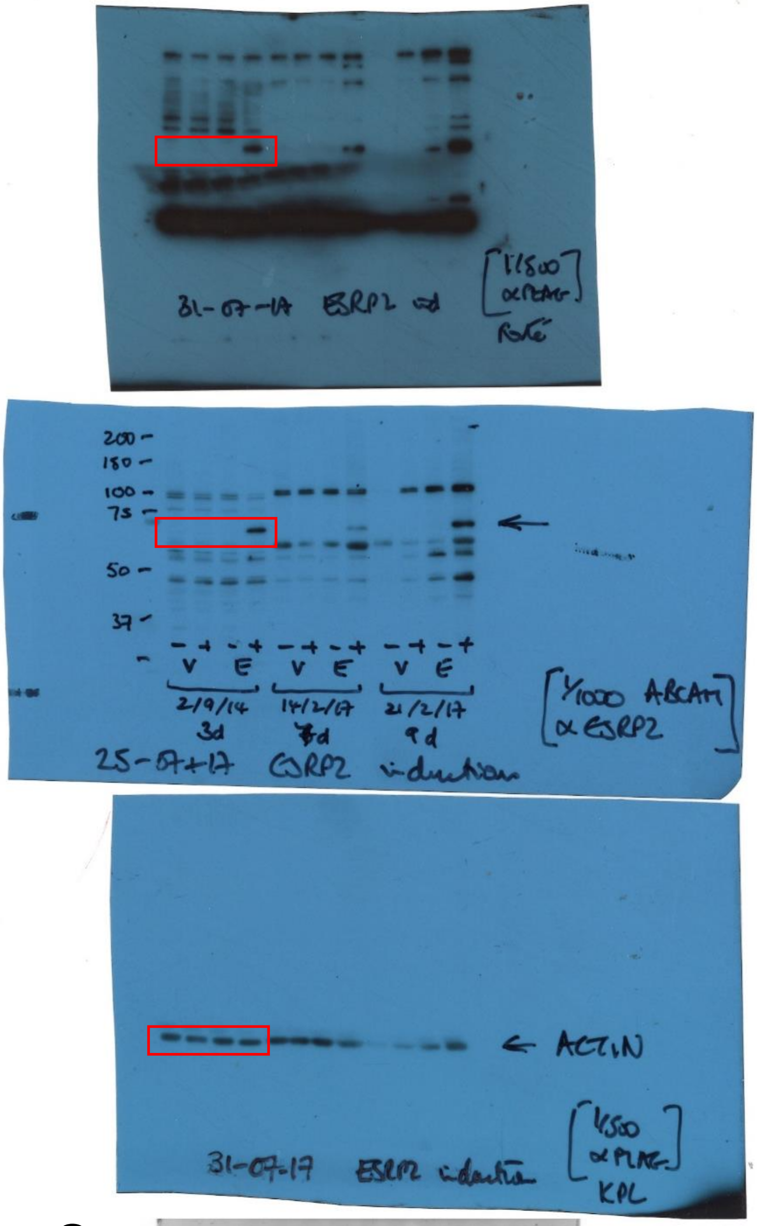


C

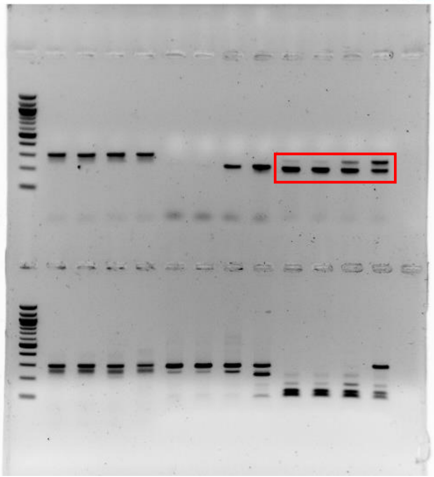

**Figure S16: Uncropped blots and gel for figure 4**
Areas used in figure 4 are outlined in red.

C

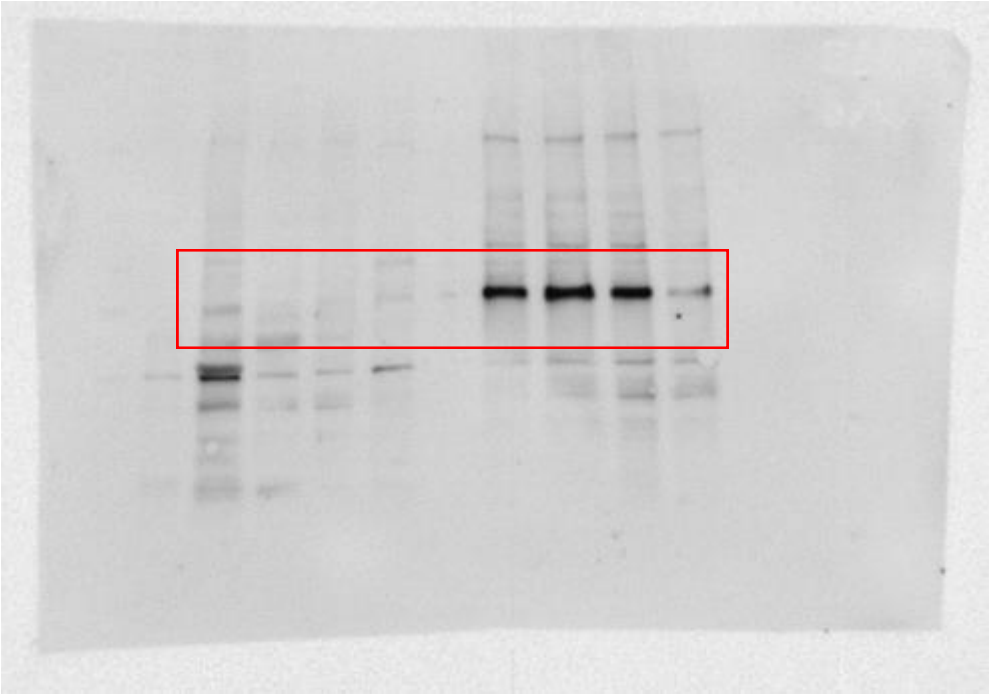

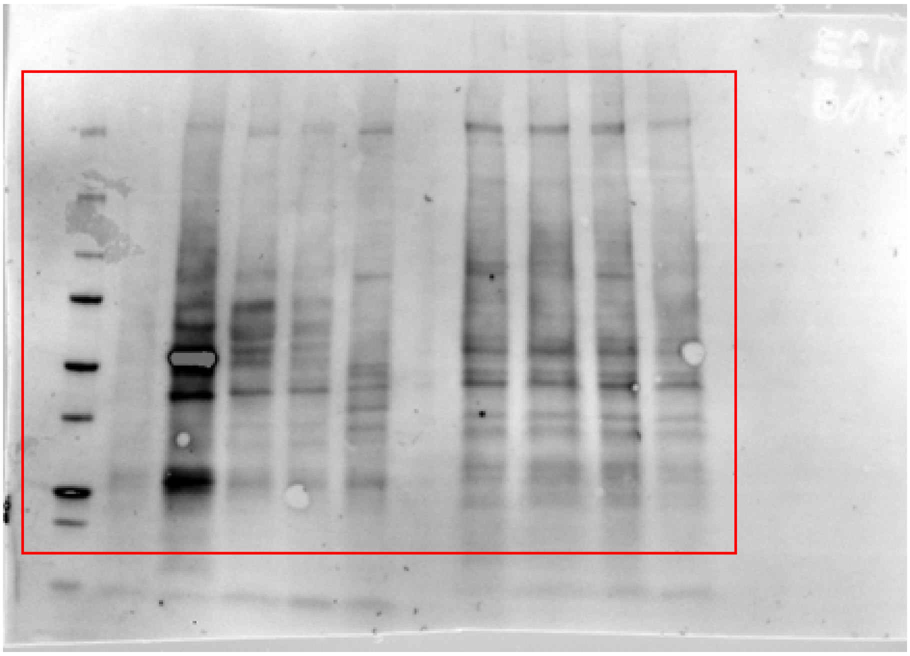

**Figure S17: Uncropped blots and gel for figure 5**
Areas used in figure 5 are outlined in red.

D

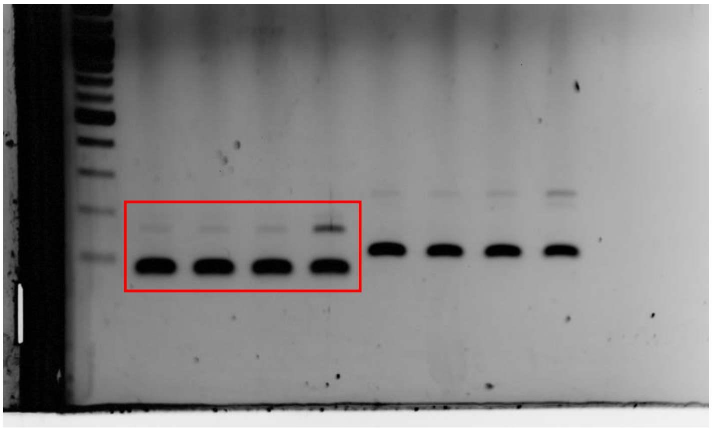

E

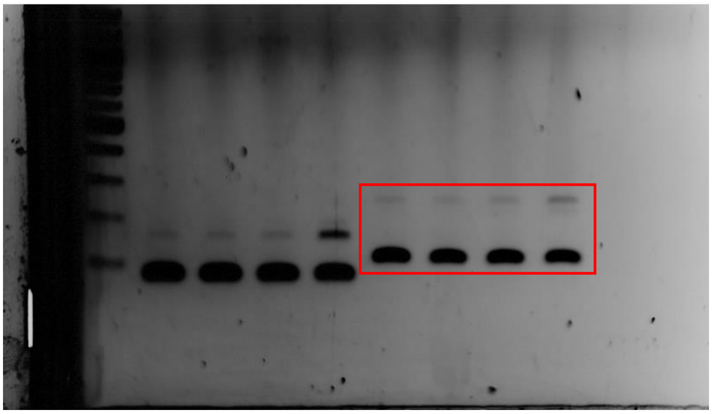

F

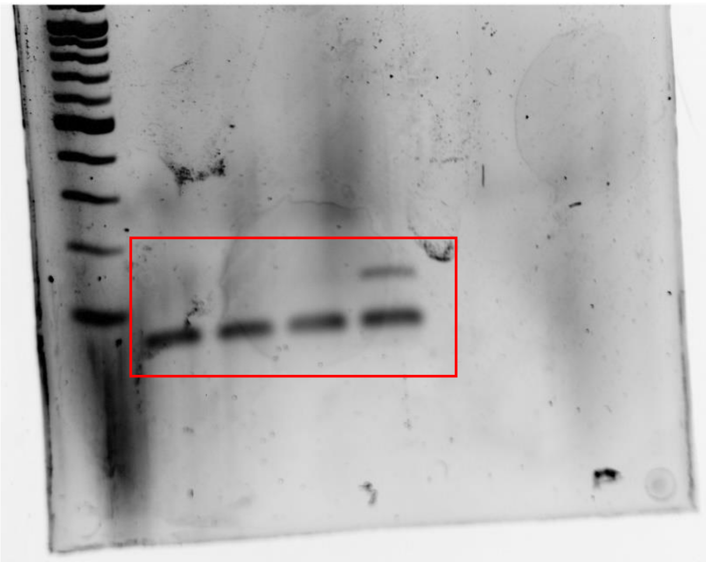

**Figure S18: Uncropped gels for figure 6**
Areas used in figure 6 are outlined in red.

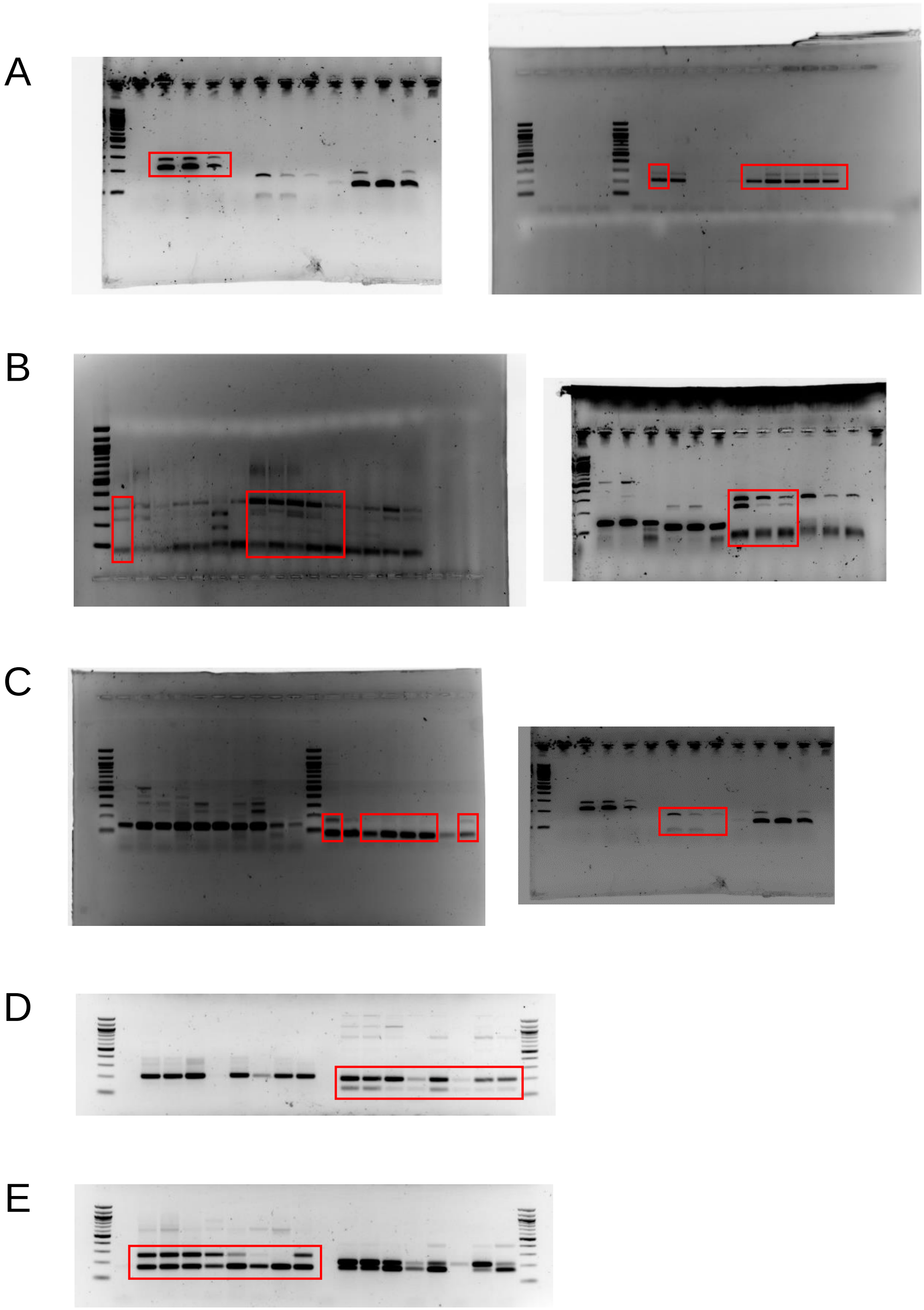


**Figure S19: Uncropped gels for figure 7**
Areas used in figure 7 are outlined in red.

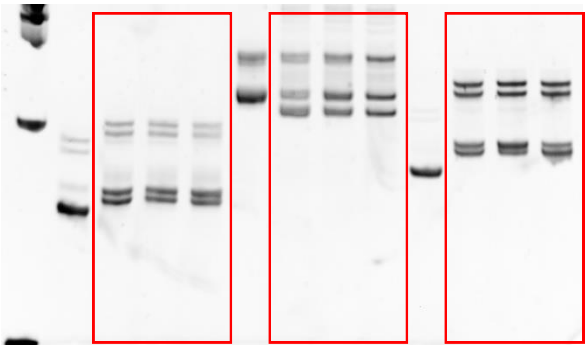

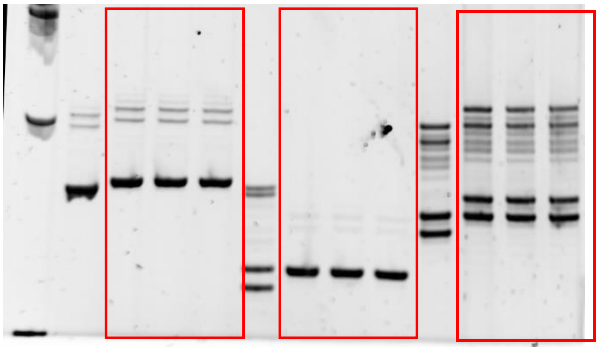

**Figure S20: Uncropped gels for figure S1B**
Areas used in figure S1B are outlined in red.

B

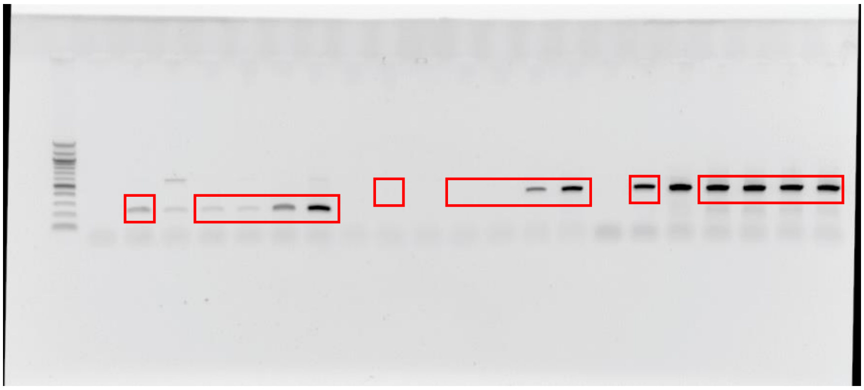

C

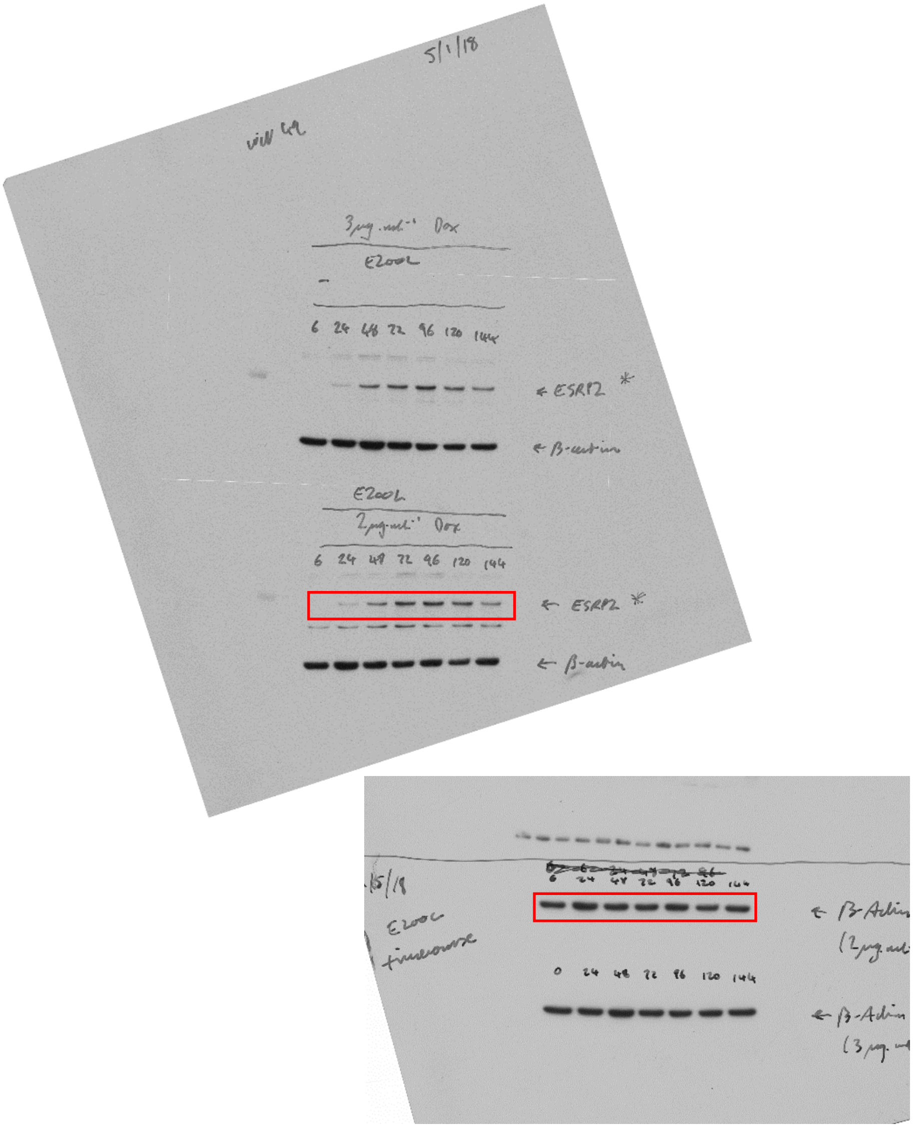


**Figure S21: Uncropped gels and blots for figure S2B and S2C**
Areas used in figure S2B and S2C are outlined in red.

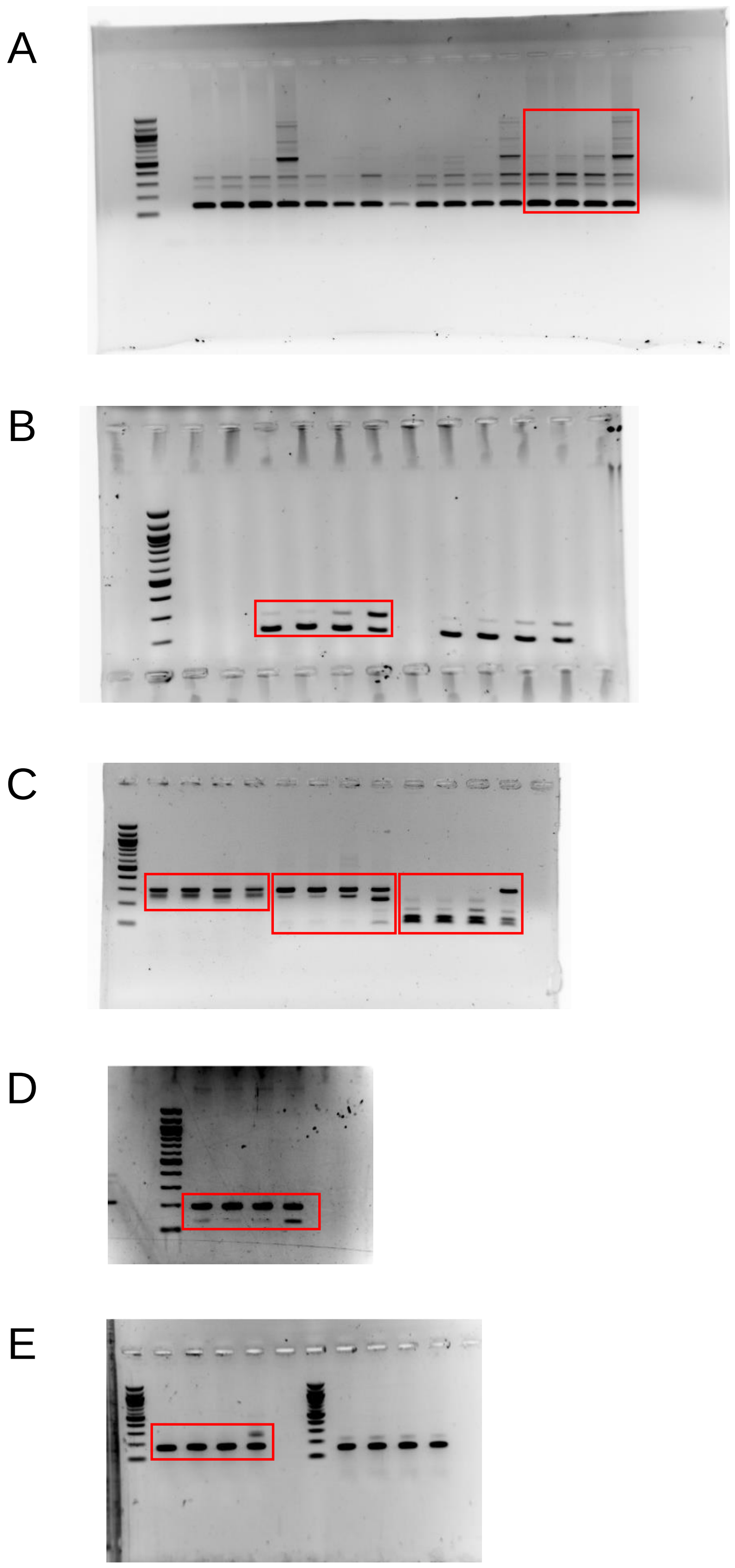


**Figure S22: Uncropped gels for figure S12**
Areas used in figure S12 are outlined in red.

A

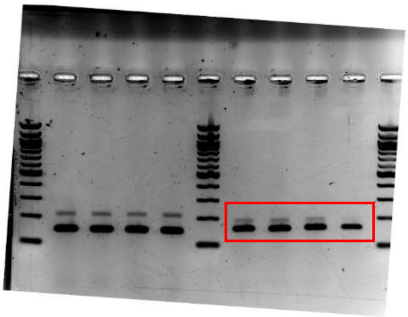

B

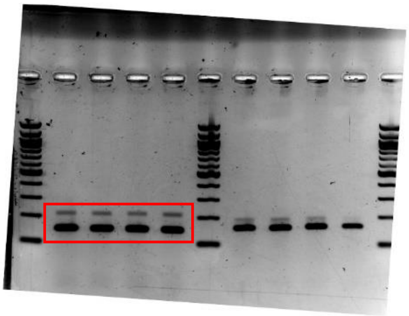

C

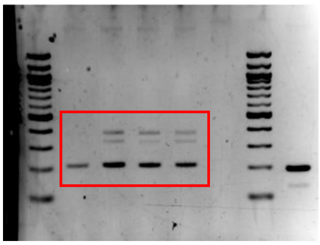

D

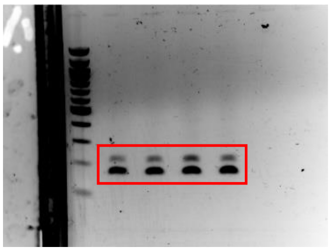

**Figure S23: Uncropped gels for figure S13**
Areas used in figure S13 are outlined in red.

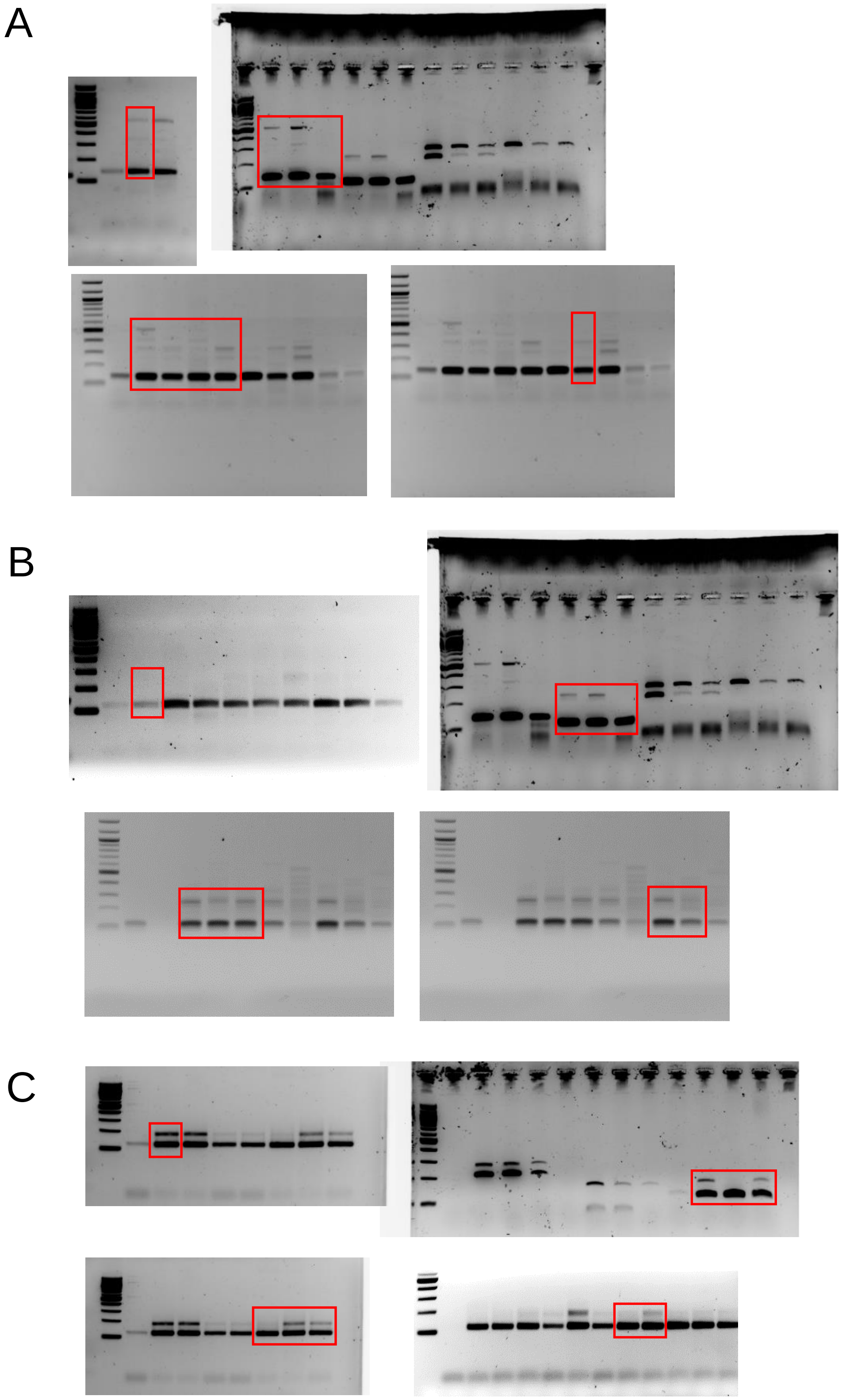


**Figure S24: Uncropped gels for figure S14**
Areas used in figure S14 are outlined in red.
